# A computational solution for bolstering reliability of epigenetic clocks: Implications for clinical trials and longitudinal tracking

**DOI:** 10.1101/2021.04.16.440205

**Authors:** Albert T. Higgins-Chen, Kyra L. Thrush, Yunzhang Wang, Pei-Lun Kuo, Meng Wang, Christopher J. Minteer, Ann Zenobia Moore, Stefania Bandinelli, Christiaan H. Vinkers, Eric Vermetten, Bart P.F. Rutten, Elbert Geuze, Cynthia Okhuijsen-Pfeifer, Marte Z. van der Horst, Stefanie Schreiter, Stefan Gutwinski, Jurjen J. Luykx, Luigi Ferrucci, Eileen M. Crimmins, Marco P. Boks, Sara Hägg, Tina T. Hu-Seliger, Morgan E. Levine

## Abstract

Epigenetic clocks are widely used aging biomarkers calculated from DNA methylation data. Unfortunately, measurements for individual CpGs can be surprisingly unreliable due to technical noise, and this may limit the utility of epigenetic clocks. We report that noise produces deviations up to 3 to 9 years between technical replicates for six major epigenetic clocks. The elimination of low-reliability CpGs does not ameliorate this issue. Here, we present a novel computational multi-step solution to address this noise, involving performing principal component analysis on the CpG-level data followed by biological age prediction using principal components as input. This method extracts shared systematic variation in DNAm while minimizing random noise from individual CpGs. Our novel principal-component versions of six clocks show agreement between most technical replicates within 0 to 1.5 years, equivalent or improved prediction of outcomes, and more stable trajectories in longitudinal studies and cell culture. This method entails only one additional step compared to traditional clocks, does not require prior knowledge of CpG reliabilities, and can improve the reliability of any existing or future epigenetic biomarker. The high reliability of principal component-based epigenetic clocks will make them particularly useful for applications in personalized medicine and clinical trials evaluating novel aging interventions.

## Introduction

Biological age estimation has been pursued to study the aging process, predict individual risk of age-related disease, and evaluate the efficacy of aging interventions (Jylhävä et al. 2017). A variety of epigenetic clocks based on DNA methylation have been developed to predict biological age and are among the most studied biomarkers of aging (Bell et al. 2019; Jylhävä et al. 2017; Horvath & Raj 2018). In humans, these clocks primarily utilize methylation arrays, particularly Illumina Infinium BeadChips that measure hundreds of thousands of specific CpG methylation sites. Most existing clocks were trained by applying supervised machine learning techniques to select a subset of CpG sites (usually a few hundred) for a weighted linear prediction model of age or aging phenotypes such as mortality risk. Thus, the prediction value of epigenetic clocks depends on the aggregate prediction value of individual CpGs.

Alas, previous studies have shown that the majority of individual CpGs are unreliable, yielding surprisingly variable methylation values when the same biological specimens are measured multiple times due to technical variance inherent to the array (Sugden et al. 2020; Logue et al. 2017; Bose et al. 2014). In many cases, technical variance exceeds the biological variance for the DNAm levels at individual CpGs, as quantified by the intraclass correlation coefficient (ICC) metric. Poor reliability has been found for consistent subsets of CpGs across multiple studies, suggesting DNAm unreliability is replicable and systematic. Technical variation can stem from sample preparation, the number of beads per CpG on each chip, probe hybridization issues (cross-reactivity, repetitive DNA, or genetic variation at probe binding sites), probe chemistry (differences between Infinium type I and type II probes), batch effects, and platform differences (e.g., 450K vs. EPIC arrays) (Naeem et al. 2014; Bose et al. 2014; Pidsley et al. 2016; Logue et al. 2017). Various processing methods may reduce technical variance, including normalization, batch correction based on control probes, stringent detection thresholds to address background noise, or limiting the analysis to high-quality probes (Lehne et al. 2015; Morris & Beck 2015; Naeem et al. 2014). However, significant unreliability remains post-processing (Sugden et al. 2020; Lehne et al. 2015).

Ultimately, there is a signal (biological variation) vs. noise (technical variation) problem for epigenetic clocks. CpGs with high biological variance tend to have higher reliability (Sugden et al. 2020; Bose et al. 2014). Sugden and colleagues showed technical variance is large enough relative to biological variance to cause wide-ranging consequences for epigenetics studies. In particular, they noted that the widely used Horvath, Hannum, and PhenoAge clocks contain many unreliable CpGs. However, it was difficult to determine the implications of these unreliable clock CpGs, given that the reported CpG reliability values were calculated from a cohort where all participants were 18 years old, limiting the biological variance one could observe. Thus, it is unknown how age-related variance quantitatively compares to technical variance at clock CpGs, given that aging has widespread effects on DNA methylation (Horvath & Raj 2018; Liu et al. 2020). The aggregate effects of the hundreds of CpGs (each weighted differently) that compose epigenetic clocks on overall clock reliability have also not been characterized. It is possible, for example, that the machine learning techniques used to train epigenetic clocks penalize noisy CpGs or select CpGs that cancel out noise from one another. However, a study examining 12 samples indicated that the Horvath multi-tissue predictor can deviate between technical replicates by a median of 3 years and up to a maximum of 8 years, and these deviations remain high regardless of preprocessing method (McEwen et al. 2018).

The threat of technical noise has major implications for utilizing epigenetic clocks in basic and translational research. For the vast majority of epigenetic aging studies in which epigenetic age is estimated cross-sectionally, noise could lead to mistaken measurements for a substantial number of individuals. There is also great interest in short-term longitudinal measurements of individuals’ biological ages, including for clinical trials and personalized medicine that aim to improve health by modifying biological age. Such studies may be particularly vulnerable to technical factors. For example, if a treatment is capable of causing a 2-year reduction in epigenetic age relative to placebo, technical variation of up to 8 years may obfuscate this effect.

In the present manuscript, we describe how technical variation leads to significant deviations between replicates for many epigenetic clocks. To address this critical issue, we provide a novel computational solution that extracts the shared aging signal across many CpGs while minimizing noise from individual CpGs.

## Results

### Poor-reliability CpGs reduce the reliability of epigenetic age prediction

To investigate the impact of low-reliability CpGs on epigenetic clock predictions, we examined the publicly available dataset GSE55763 (Lehne et al. 2015) comprising 36 whole blood samples with 2 technical replicates each and an age range of 37.3 to 74.6. The data was processed to eliminate systematic technical bias between batches (see Materials and Methods). In an ideal scenario, the same sample measured twice using methylation arrays would yield the same age prediction. Deviations from this ideal can be quantified using the intraclass correlation coefficient (ICC), a descriptive statistic of the measurement agreement for multiple measurements of the same sample (within-sample variation) relative to other samples (between-sample variation) (Koo and Li 2016). Biological between-sample variance can correspond to age, sex, smoking, genetics, cell composition, and other factors. Technical within-sample variance can arise from sample preparation, hybridization to the methylation array, scanning, data processing, and stochastic factors. Using a dataset with a wide age range is critical to determine the relative degrees of age-related variance compared to technical variance.

First, we calculated ICCs for 1,273 individual CpGs that are part of five existing clocks— the Horvath multi-tissue predictor (Horvath1), the Horvath skin-and-blood clock (Horvath2), the Hannum blood clock (Hannum), the Levine DNAmPhenoAge clock (PhenoAge), or the Lu telomere length predictor (DNAmTL) (Table S1) (Horvath 2013; Horvath et al. 2018; Hannum et al. 2013; Levine et al. 2018; Lu, Seeboth, et al. 2019). 31.6% of clock CpGs have reliabilities characterized as poor (ICC <0.5), 21.5% as moderate (0.5 <= ICC < 0.75), 29.9% as good (0.75 <= ICC < 0.9), and 17.0% as excellent (ICC >= 0.9) (Figures 1A, S1, Table S2). Low-reliability clock CpGs tend to have more extreme values (near 0 or 1) and lower variance (Figures 1B-C), consistent with prior genome-wide analyses (Sugden et al. 2020; Bose et al. 2014). CpGs with strong associations with mortality (hazard ratios after adjusting for age and sex) or with chronological age tended to have higher ICCs (Figure 1D). However, age and mortality associations are artificially depressed by high technical noise and thus low-ICC CpGs likely still contain useful information. CpGs within individual clocks largely showed the same patterns (Figures S2-S8).

**Figure 1.**
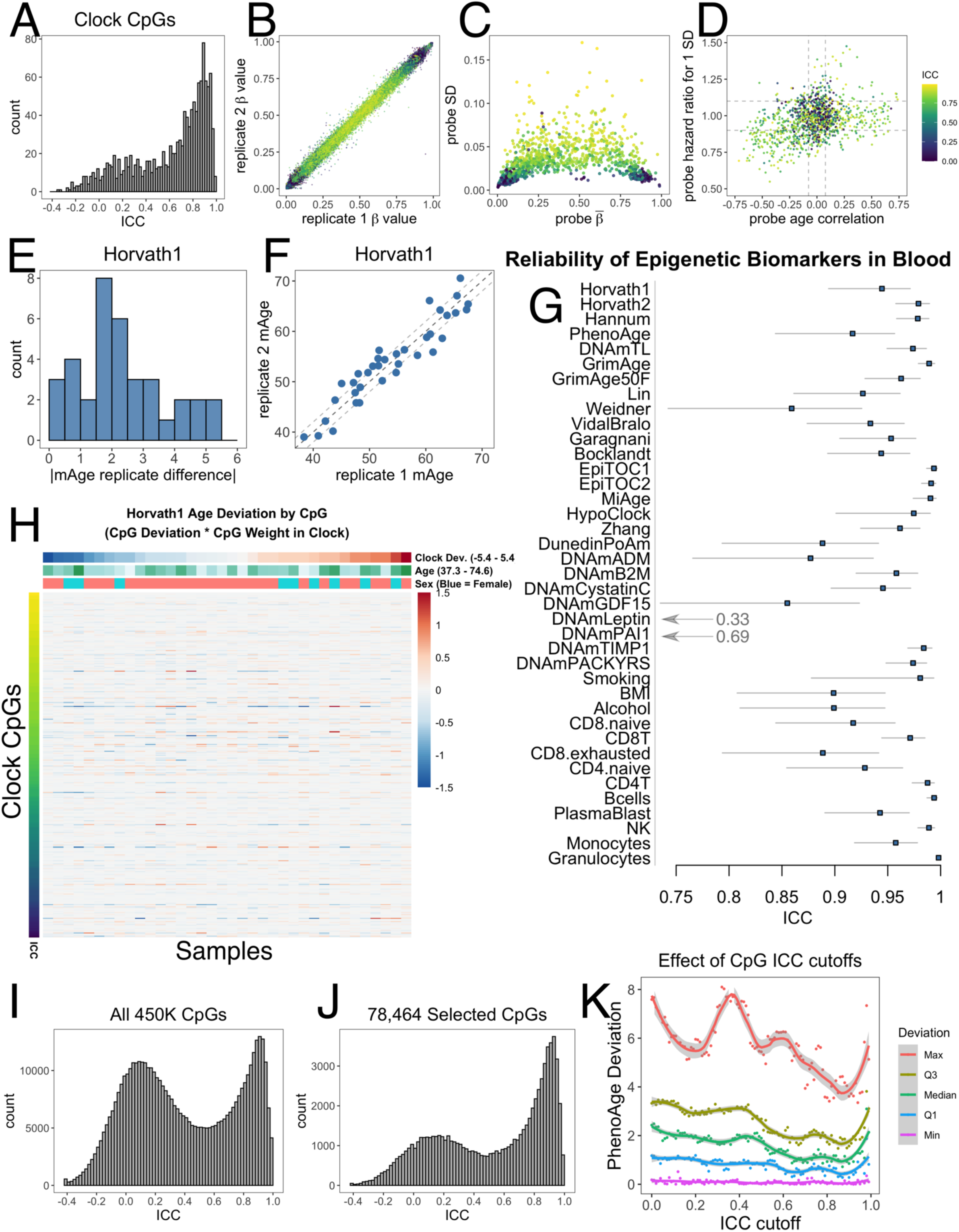
Poor-reliability CpGs reduce the reliability of epigenetic age prediction. A) Intraclass correlation coefficients (ICCs) for 1,273 CpGs in the Horvath1, Horvath2, Hannum, PhenoAge, or DNAmTL clocks, analyzed in 36 pairs of technical replicates in blood (GSE55763). B) Clock CpG ICCs versus beta values for all samples. Each point corresponds to one pair of replicates for one CpG. C-D) Comparisons of clock CpG ICCs to CpG mean beta value, standard deviation, age correlation (in GSE40279), and mortality hazard ratio (in the Framingham Heart Study, after adjusting for age and sex). Each point corresponds to one CpG. E-F) Deviations between replicates in Horvath1. Each point corresponds to one sample. G) ICCs for epigenetic biomarkers (raw score not adjusted for age). H) Contribution of each CpG to the total age deviation between replicates for Horvath1. I) ICCs for all 450K CpGs. J) ICCs for selected 78,464 CpGs present on 450K and EPIC arrays as well as training, test, and validation data sets. This number was small enough for repeated experimentation with methods to improve reliability. K) Filtering CpGs by ICC cutoffs prior to training a predictor of PhenoAge only modestly improves reliability.

We then investigated the reliability of overall epigenetic clock scores and found all the clocks demonstrated substantial biological age discrepancies between technical replicates. The widely used Horvath1 multi-tissue clock (Horvath 2013) shows a median difference of 2.1 years between replicates, and a maximum deviation of 5.4 years (Figure 1E-F). For other clocks, median deviation ranges from 0.9 to 2.4 years and maximum difference ranges from 4.5 to 8.6 years, depending on the clock (Figure S9-S10). Clock ICCs range from 0.917 to 0.979, with the Horvath1 multi-tissue clock exhibiting an ICC of 0.945 (Figure 1G; Table S3). Epigenetic age acceleration (i.e. after adjusting for chronological age) is commonly used in models of aging outcomes and is arguably the measure one would care about most for intervention studies. Age acceleration has lower ICCs because of reduced biological variance, with ICCs ranging from 0.755 to 0.948 (Horvath1 ICC = 0.817) (Table S4). The discrepancies of different clocks are mostly uncorrelated with each other and with age and sex (Figure S11), consistent with their origin in noise. There was little to no systematic batch effect (i.e. no bias in the direction of deviation) for any clock except for Horvath2, which showed a small effect (0.6 years mean difference between batches, p=0.0486) (Table S5). All samples showed substantial deviations in at least one clock (Table S5), demonstrating unreliability is not confined to specific samples.

We also assessed the reliability of GrimAge (Lu, Quach, et al. 2019), a unique case because chronological age and sex are required for its calculation and CpG identities are not published. GrimAge showed better reliability than other clocks (median deviation between replicates = 0.9 years; maximum deviation = 2.4 years; GrimAge ICC = 0.989; GrimAgeAccel ICC = 0.960; Figure 1G; Tables S3-4). The ICC of GrimAge components was lower, ranging from 0.329 to 0.984. Since age and sex are the same for technical replicates but different between biological samples, we isolated the variation attributed to DNAm by re-calculating GrimAge after artificially setting age to 50 and sex to female for all samples (GrimAge50F). The ICC decreased from 0.989 to 0.963, below that of Horvath1, Horvath2, and Hannum. Thus, DNAm unreliability also affects GrimAge, when considering its values independent of age and sex.

We further calculated ICCs for various epigenetic biomarkers (Figure 1G, Tables S3-4), including additional aging clocks, estimated cell proportions, and predictors of smoking, alcohol and BMI (Horvath 2013; Horvath et al. 2018; Hannum et al. 2013; Levine et al. 2018; Lu, Seeboth, et al. 2019; Lu, Quach, et al. 2019; Lin & Wagner 2015; Weidner et al. 2014; Vidal-Bralo et al. 2016; Garagnani et al. 2012; Bocklandt et al. 2011; Teschendorff 2020; Youn & Wang 2018; Belsky et al. 2020; McCartney et al. 2018; Houseman et al. 2012; Zhang et al. 2017). Mitotic clocks and granulocytes had particularly high ICCs, but otherwise unreliability issues affect nearly all epigenetic biomarkers.

To better understand how technical variation in individual CpGs contributes to the technical variation in each clock overall, we multiplied the CpG differences between technical replicates by the CpG clock weights. We plotted the resulting values for all CpGs and all samples in heatmaps (Figures 1H, S12). Globally, different sets of CpGs contribute to technical variation in different samples, consistent with noise. These effects can be quite large--for each clock, some individual CpGs each contribute more than 1.5 years to the total discrepancy between specific pairs of replicates. The direction of effect for these CpGs is nonuniform, suggesting these are not batch effects that can be easily corrected for. Deviations are distributed throughout the age range and occur in both sexes. Even within samples that do not show overall clock deviations between replicates, there is significant noise from individual CpGs but their effects happen to cancel each other out. CpGs contributing a relatively large amount to total noise are distributed throughout the range of ICC values, suggesting this noise may be difficult to filter out. These results demonstrate that reliability issues are not limited to specific samples or a small subset of CpGs. Overall, these findings confirm that the poor reliability of individual CpGs is a significant problem for epigenetic age prediction.

### Filtering CpGs by ICC only modestly improves reliability

We turned our attention to a broader set of CpGs with the goal of developing clocks with superior reliability, without sacrificing validity. We selected 78,464 CpGs (Table S6) that were present in a wide range of DNAm datasets for training, testing, and validation obtained on either the EPIC or 450K Illumina arrays. These datasets were curated to represent many different tissues and include some with technical replicates or longitudinal follow-up (Table S7). This set of CpGs was small enough for repeated computations to systematically explore different methods to improve reliability, but still larger than the set of 27K CpGs used to train the original Horvath1 and PhenoAge clocks. The reliability distribution of this subset is similar to that of existing clocks and has fewer poor-reliability CpGs compared to the set of all 450K CpGs (Figure 1A, I, J; Table S6). We found a bimodal distribution of CpG reliabilities consistent with prior studies (Bose et al. 2014; Dugue et al. 2016), and our ICCs were well-correlated with previously published ICC values from 3 prior studies (Figure S13; Table S6) (Bose et al. 2014; Logue et al. 2017; Sugden et al. 2020). Poor-reliability CpGs still show age and mortality correlations in independent datasets, albeit lower than high-reliability CpGs (Figure S14). We note this does not necessarily mean that poor-reliability CpGs have less information about aging; instead, technical noise may dilute the signal.

The most intuitive solution to address DNAm reliability is to filter out unreliable CpGs based on ICC values, then re-train epigenetic clocks. Thus, we systematically tested the effects of various ICC cutoffs when considering which CpGs to include in supervised learning for phenotypic age prediction in the original DNAmPhenoAge training data (InCHIANTI). We chose to demonstrate this using DNAmPhenoAge given its low ICC, yet high mortality prediction, with the goal of testing whether ICC could be improved without jeopardizing the latter. Reliability improved modestly when training clocks with a subset of CpGs with higher ICC cutoffs (Figure 1K, Figure S15, Table S8), whereas no improvement occurred when taking an equivalent number of random CpGs (Figure S16). Discarding all poor-reliability CpGs with ICC < 0.5 still results in maximum deviations of 6 years. An optimum cutoff occurs at an ICC cutoff of 0.9 (after discarding 80% of CpGs), but maximum deviations continue to be 4 years, the third quartile 2 years, and median 1 year. Unfortunately, we also found that when one progressively drops CpGs using cutoffs above 0.9, mortality prediction decreases sharply and between-replicate deviations increase, possibly because there are now insufficient numbers of CpGs available for training (less than 15,000 CpGs) (Figure S15).

We also experimented with other methods such as introducing a penalty factor for each CpG inversely proportional to ICC into elastic net regression, as well as training clocks using M-values or winsorized beta-values (Table S9). However, we did not find these substantially improved clock reliability.

Thus, straightforward filtering approaches to poor-reliability CpGs only modestly improves clock reliability. Indeed, the existing Hannum clock is already mostly composed of CpGs with high age correlations and almost no poor-reliability CpGs (Figures S6-S8), but still shows deviations between technical replicates up to 5 years (Figure S10) with many individual CpGs contributing deviations of up to 1.5 years each (Figure S12). Filtering requires *a priori* knowledge of CpG reliabilities, which is often not known for a given tissue or sample population and thus this approach would not be easily generalizable. Filtering out more than 80% of CpGs deemed as “unreliable” will likely eliminate relevant information about aging in non-blood tissues, adiposity, smoking, alcohol, and other age- and mortality-related phenotypes. Overall, we concluded that epigenetic clocks trained on individual CpGs come with inherent noise that is not easily discarded.

### Epigenetic clocks trained from principal components are highly reliable

Many CpGs tend to change together with age in a multicollinear manner, including far more CpGs than those found in existing clocks (Liu et al. 2020; Higgins-Chen et al. 2021; Horvath & Raj 2018). Elastic net regression, commonly used to build clocks, uses model penalties to select a limited number of CpGs to represent a set of collinear CpGs while avoiding overfitting. However, by virtue of incorporating methylation information from individual CpGs, these models retain much of the technical noise that exists in DNA methylation array technology (Figure 1). We hypothesized that principal component analysis (PCA) could extract the covariance between multicollinear CpGs, including age-related covariance. At the same time, since technical noise does not appear to be significantly correlated across CpGs after adequate preprocessing and batch correction (Figure 1H), the principal components may remain largely agnostic to technical noise.

PCs were estimated from the 78,464 CpGs in the datasets used to train epigenetic clocks (Table S7). Since each clock was trained using different data, we calculated a separate set of PCs for each clock. For all instances, the ICCs of PCs were much higher than those of individual CpGs, despite being derived from those same CpGs (Figure 2A). We then applied elastic net regression to retrain 6 epigenetic clocks from PCs (Figure 2B), and projected test datasets onto the training PCA space which allowed us to calculate and then validate the new clocks in independent data. We found it is possible to predict either the original outcome variable (e.g. age, phenotypic age), or the original CpG-based epigenetic clock score from PCs. This latter option is useful in cases where not all of the original training data is available (see Methods), in order to maintain consistency with existing studies utilizing CpG-based clocks. For example, the Horvath1 multi-tissue predictor was trained using both 27K and 450K data, but our set of 78,464 CpGs must be obtained on either 450K or EPIC, so we found replacement datasets for the 27K data. We ultimately chose to train PC clock proxies of the original CpG clock versions of Horvath1, Horvath2, Hannum, DNAmTL, and GrimAge. In contrast, PCPhenoAge was trained directly on phenotypic age based on clinical biomarkers rather than DNAm (Levine et al. 2018), but still correlates well with DNAmPhenoAge in test data (Figure 2F). These PC-based clocks showed high correlations with the original CpG versions within both training and test datasets (Figures 2C-H), with the elastic net cross-validation procedure selecting anywhere from 120 to 651 PCs.

**Figure 2.**
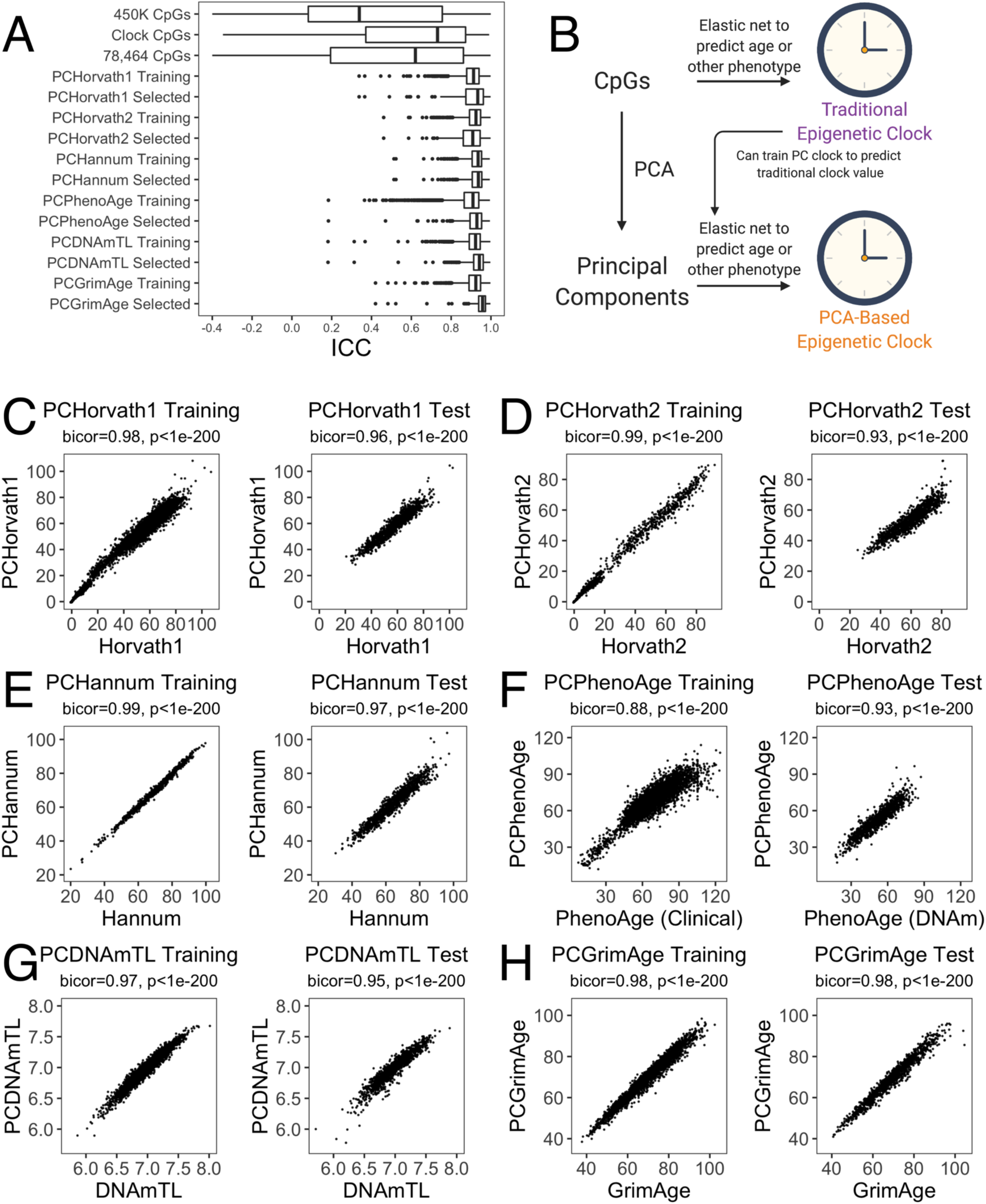
Epigenetic clocks trained from principal components. A) ICC distributions for PCs in test data compared to CpGs. B) Strategy for training PC clocks compared to traditional epigenetic clocks. Image created with Biorender.com. C-H) Correlations between the original clocks and their PC clock proxies in both training and test data. Test data shown is the Framingham Heart Study methylation data for all clocks, using samples that were not used to train PCDNAmTL or PCGrimAge.

Even though our models were trained naïve to any information about technical replicates or ICCs, the PC-based clocks showed greatly improved agreement between technical replicates (Figure 3A-F). Most replicates (90% or more) showed agreement within 1-1.5 years. The median deviation ranged from 0.3 to 0.8 years (improvement from 0.9-2.4 years for CpG clocks). All ICCs improved substantially, with all PC clocks having ICC greater than 0.99 for raw clock score and 0.97 for age acceleration (Figure 3G-H).

**Figure 3.**
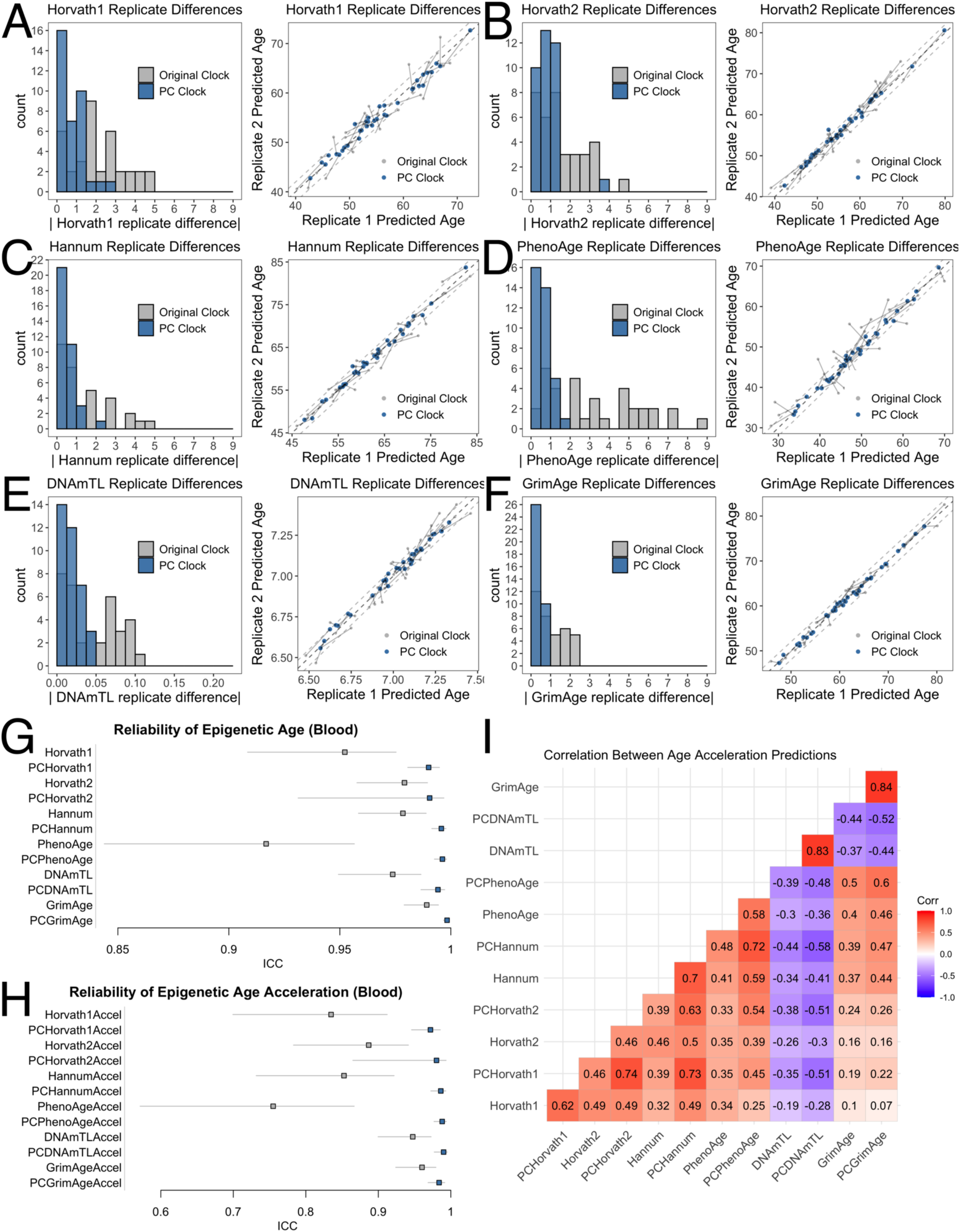
Epigenetic clocks trained from principal components are highly reliable. A-F) Agreement between technical replicates in blood test data (GSE55763). G-H) ICCs for (G) epigenetic clock scores without residualization and (H) epigenetic age acceleration in GSE55763. I) Correlation between age acceleration values for original and PC clocks in FHS blood test data.

The most dramatic improvement was in PhenoAge. CpG-trained PhenoAge has a median deviation of 2.4 years, 3^rd^ quartile of 5 years, and maximum of 8.6 years. In contrast, PCPhenoAge has a median deviation of 0.6 years, 3^rd^ quartile of 0.9 years, and maximum of 1.6 years. For this version of the clock, we added methylation and phenotypic age data from the Health and Retirement Study (HRS) (Crimmins et al. 2021). However, the same improvement in reliability occurred regardless of training on only the original InCHIANTI dataset (Table S10), or on the combined InCHIANTI and HRS dataset (Figure 3D). Notably, this improvement was far superior to filtering out even 80% of the lowest ICC CpGs (Figure 1K).

The age acceleration values were also highly correlated between PC clocks and their CpG counterparts (Figure 3I). Also, the correlations between different PC clocks was stronger than between CpG clocks. This may be partly due to the shared set of CpGs used to train PCs, or due to the reduction of noise that would have biased correlations towards the null. Correlations between PC clocks and CpG clocks tended to be stronger compared to correlations between CpG clocks and CpG clocks, consistent with a reduction of noise.

We tested if incorporating reliability information into the training method for PC clocks further increased clock reliability (Table S11), given that modest improvement occurred with CpG filtering in Figure 2. However, filtering out poor-reliability CpGs prior to PC training or pre-selecting PCs based on ICC or variance explained does not improve reliability. Re-introducing high-reliability CpGs to PCs immediately reduces clock reliability even when limiting to CpGs with ICCs greater than 0.99.

In fact, when we examined the loadings for CpGs on the PCs, we found poor-reliability CpGs contribute substantially to the overall PC clocks (Figure S17), consistent with the hypothesis that PCA can extract the denoised age signal from poor-reliability CpGs by leveraging the covariance between CpGs. This is further evidence that low-reliability CpGs likely contain important information about aging and should not simply be discarded.

### PC clocks allow for correction of systematic offsets in epigenetic age between batches

Batch correction is a standard preprocessing step during DNA methylation analysis (Morris & Beck 2015). However, there is a balance between adequate batch correction and over-correction that can lead to false positive and false negative results (Zindler et al. 2020). In many datasets there may be residual batch effects after correction that affect epigenetic clock predictions. We would not expect that the PC clock methodology, by itself, would resolve this issue because batch effects influence numerous CpGs simultaneously. However, because within-batch technical replicates show lower variance using PC clocks, we hypothesized that between-batch variation (such as systematic offsets in epigenetic age) should be easier to detect and correct for. To test this, we examined blood from each individual with age range 26-68 on the EPIC array (Figure 4). For each individual, we collected 3 blood samples simultaneously, ran each sample in a separate batch with 3 technical replicates each, and scanned each batch twice, for a total of 18 replicates per individual (3*3*2). We detected systematic offsets in the clocks based on batch (e.g. Horvath1 was reduced by 3 years in batch 3 compared to batch 1, while PCHorvath1 was reduced by 4 years) (Table S12). Correcting for batch in a linear model led to strong agreement between replicates regardless of batch for PC clocks, but not for CpG clocks.

**Figure 4.**
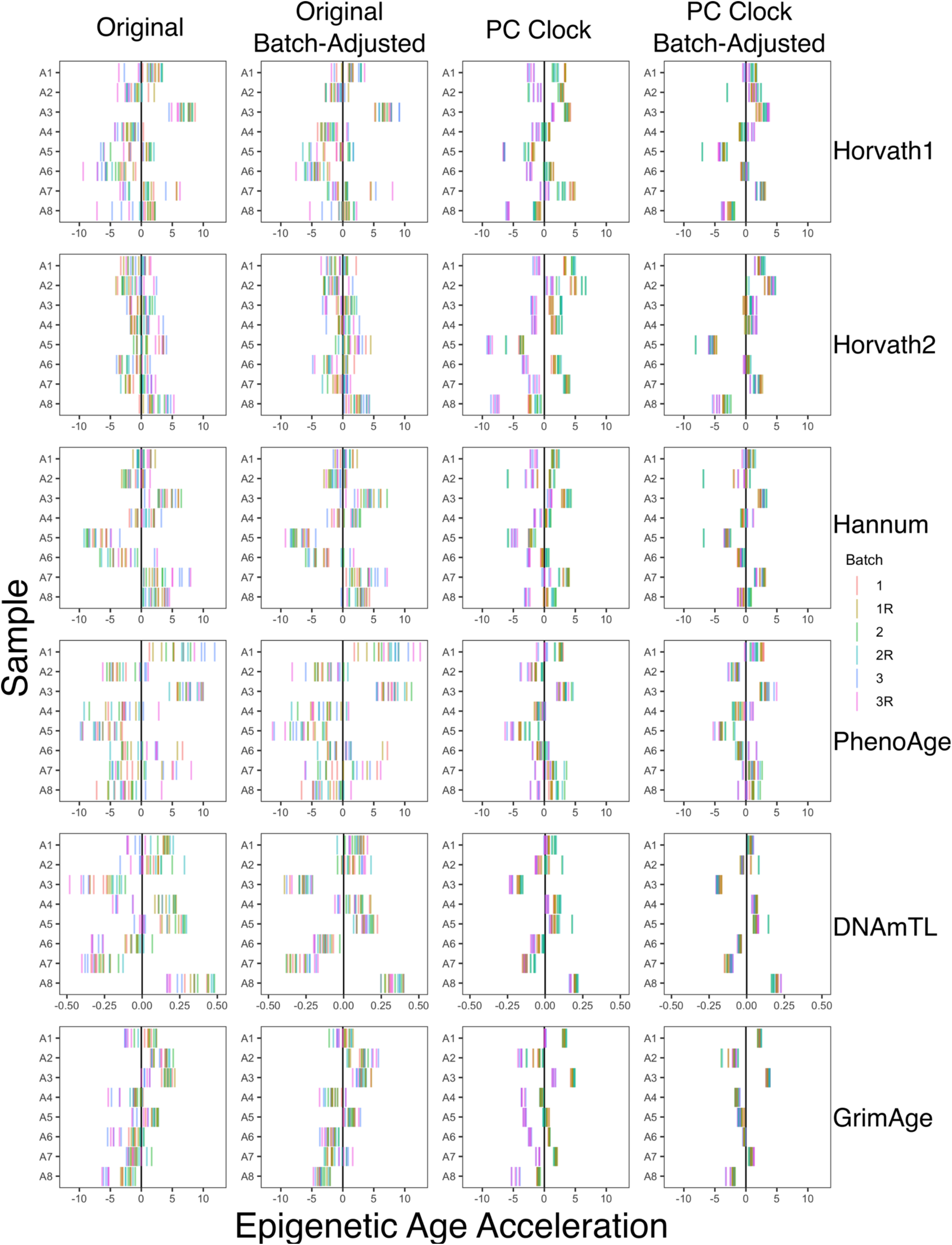
PC clocks allow for correction of systematic offsets in epigenetic age between batches. The original clocks and corresponding PC clocks were calculated for 8 blood samples with 18 technical replicates each (3 batches, 3 replicates per batch, 2 scans per batch). Batch correction was performed using a linear model using batch as a categorical variable.

### PC clocks are reliable in saliva and brain

There is a paucity of technical replicate data in non-blood tissues. Many clocks are trained solely in blood (including Hannum, PhenoAge, DNAmTL, and GrimAge), though they often still correlate with age in other tissues (Liu et al. 2020; Horvath & Raj 2018). We tested if PC clocks show enhanced reliability in non-blood tissues. First, we measured DNAm in saliva from the same 8 individuals that we had obtained blood from, with the same design of 3 consecutive samples, 3 technical replicates each, and 2 scans. While there were again epigenetic age offsets between saliva batches similar to blood, these offsets were not consistent between samples, and therefore a linear batch correction was not possible (Figure 5A). The reason is not clear, but it is possible that saliva may change in cell composition with every consecutive sampling (e.g. changing proportions of epithelial cells vs. leukocytes), whereas blood is more consistent. Regardless, technical replicates for each sample still showed very strong agreement for PC clock age acceleration, with improved ICCs compared to the original clocks (Figure 5B). Note that PCHorvath1 and PCHorvath2 only improved marginally because they are tightly fit to chronological age, and therefore there is minimal biological variation in epigenetic age acceleration, especially for only 8 samples.

**Figure 5.**
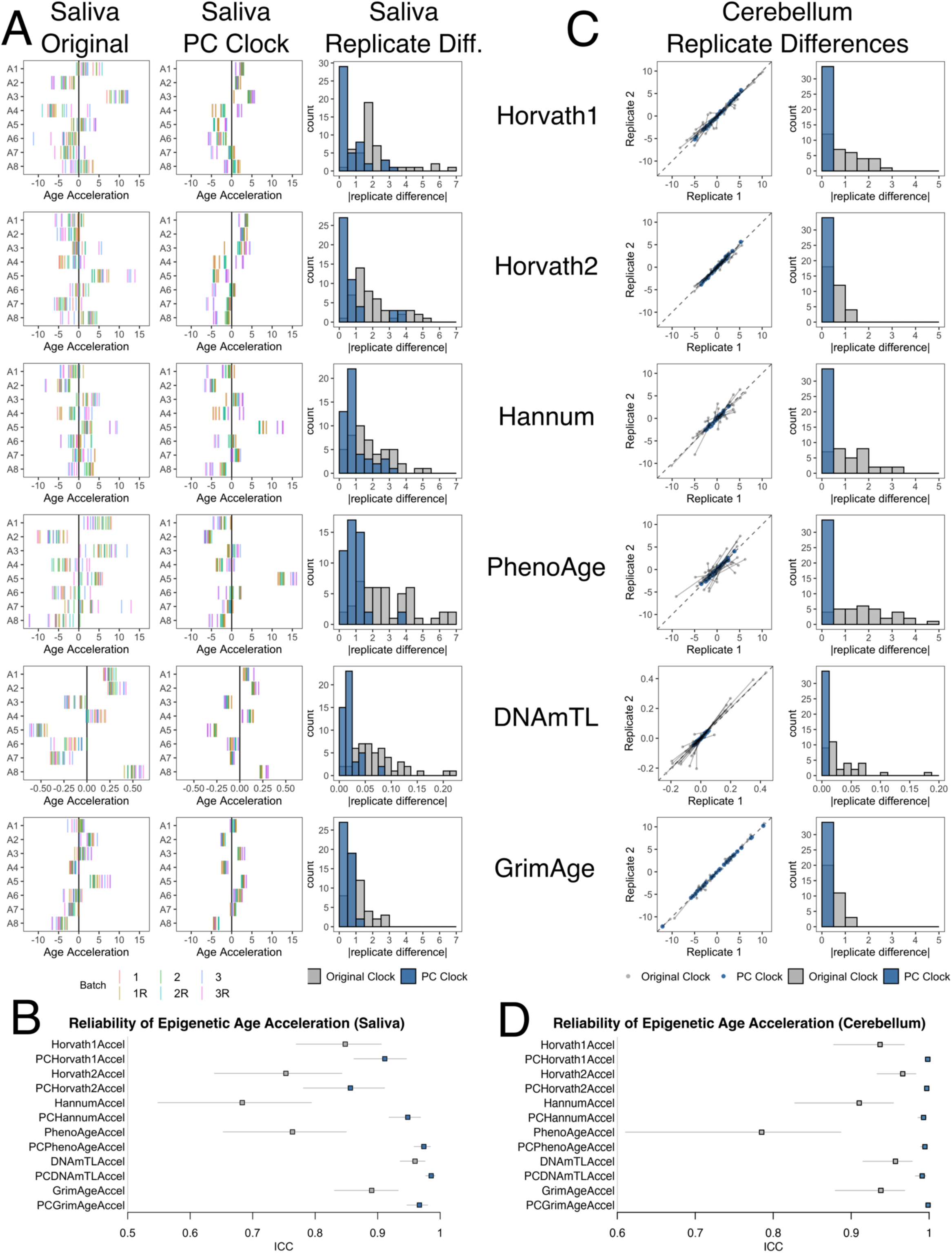
PC clocks are reliable in saliva and brain. (A) The original clocks and corresponding PC clocks were calculated for 8 saliva samples with 18 technical replicates each (3 batches, 3 replicates per batch, 2 scans per batch). (B) ICC for technical replicates in saliva, treating each batch and scan separately. (C) Agreement between technical replicates in cerebellum test data (GSE43414). Because of a systematic shift in epigenetic age between replicates, mean-centered epigenetic age values were used for both the original clocks and PC clocks. (D) ICC for technical replicates in cerebellum.

We also examined publicly available replicate data from cerebellum (GSE43414), which is known to yield lower epigenetic age predictions compared to other tissues (Horvath et al. 2015). This cohort contains data from 34 individuals, with 2 scans each. Again, we found that the two cohorts in this study display a batch shift in the original clock scores, though the PC clock scores were far more resilient against this effect (Figure S18). Thus, to provide fair comparison between the original and PC clocks, we analyzed the distance between each clock’s batch-mean centered prediction values, demonstrating that all aging clocks have an absolute disagreement of less than 0.5 years in brain tissue (Figure 5C). Although improved agreement in PCDNAmTL may be partially due to a narrower dynamic range compared to DNAmTL, there is minimal attrition of telomere length with age in the cerebellum due to high proportions of post-mitotic neurons (Demanelis et al. 2020). Thus, PCDNAmTL may be better capturing the true biological variance. We also calculated age residuals using independent linear models in each batch. The ICC values of PC clock residuals show near-perfect agreement, whereas the CpG clocks remain significantly lower in all clocks (Figure 5D).

Thus, the PC clocks are a highly reliable method for epigenetic age measurement in saliva and brain. It will be interesting to determine if future PC clocks trained in non-blood tissues show even higher reliability and greater ease of batch correction when tested in non-blood tissues.

### PC clocks preserve relevant aging and mortality signals

To test if any validity in epigenetic clocks was sacrificed in order to boost reliability, we examined associations between clocks and various sociodemographic, behavioral, and health characteristics using the Framingham Heart Study. Each of the original clocks have unique sets of associations with age-related traits and lifestyle factors, and may capture distinct aspects of aging (Levine et al. 2018; Lu, Quach, et al. 2019; Liu et al. 2020; Horvath & Raj 2018). We found that the PC versions of the clocks maintained or even exhibited improved prediction of mortality (Figure 6A). Note that GrimAge shows particularly strong mortality prediction because it was trained to predict mortality in FHS and is therefore overfit. The PC clocks also maintained associations with a wide range of other factors in the FHS cohort (Figure 6B). Overall, these preserved aging and mortality signals demonstrate that the PC clocks could be fully substituted for the original clocks in ongoing or future studies.

**Figure 6.**
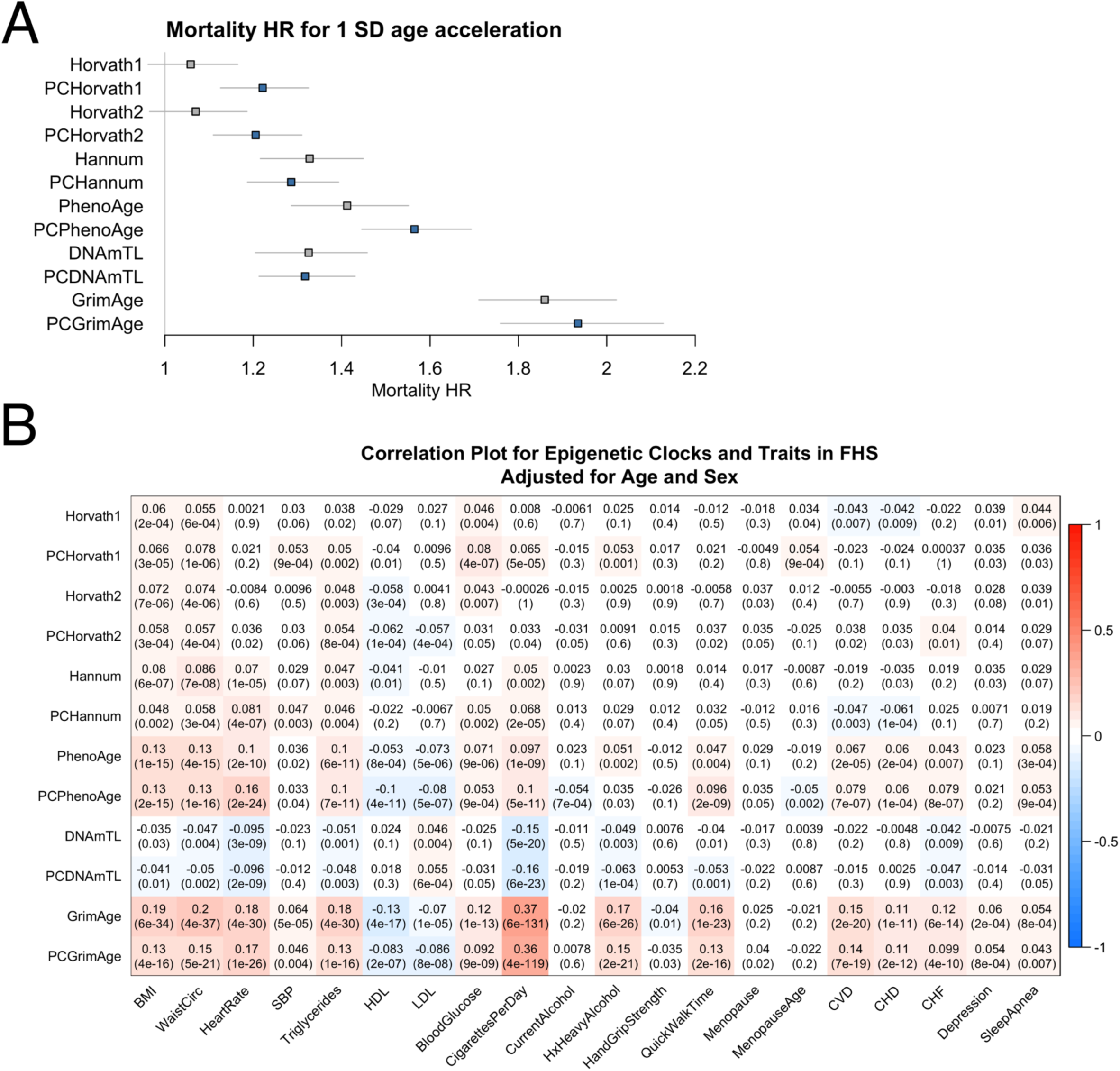
PC clocks preserve relevant aging and mortality signals. (A) Mortality hazard ratios were calculated in the Framingham Heart Study (FHS) after adjusting for chronological age and sex. (B) Correlations with various traits were calculated in FHS after adjusting for chronological age and sex. Note that GrimAge is overfit to this dataset and therefore associations are elevated compared to other clocks.

### PC clocks show improved stability in long- and short-term longitudinal data

Longitudinal studies are instrumental for studying aging as a continuous process and for assessing the utility for aging biomarkers in clinical trials and personalized medicine. Longitudinal fluctuations in epigenetic age acceleration have previously been observed (Li et al. 2020). However, if epigenetic clocks are strongly influenced by technical noise (Figure 1), this raises the concern that it may be difficult to disentangle biologically meaningful longitudinal changes (for example, those induced by lifestyle changes or a new medication) from technical variation. We hypothesized that PC clocks would show increased stability in longitudinal studies by reducing technical noise.

In longitudinal data from the Swedish Adoption Twin Study of Aging (SATSA) for 294 individuals (baseline age range 48 to 91) spanning up to 20 years of follow-up and 2 to 5 time points per person (Li et al. 2020), the original CpG clocks show age trajectories that fluctuate dramatically, deviating up to 20-30 years off the average trajectory (Figure 7A). However, the equivalent PC clocks show far less deviation. We calculated epigenetic clock slopes for every individual and found that longitudinal changes in CpG clocks were modestly correlated with those in their counterpart PC clocks. However, longitudinal changes in the original clocks are poorly correlated with each other, while changes in the PC clocks are far more tightly correlated (Figure 7B), consistent with a reduction in noise. For example, PhenoAge and GrimAge are both known predictors of mortality (Levine et al. 2018; Lu, Quach, et al. 2019), yet longitudinal change in PhenoAge is not correlated with change in GrimAge when considering the original CpG clocks. However, PCPhenoAge and PCGrimAge changes are correlated at r = 0.44. Likewise, the correlation between Horvath1 and Horvath2 longitudinal changes increases from 0.25 to 0.87 when using the PC clocks. We also calculated ICC values where higher ICC values signify reduced within-person variance, which could be composed either of technical variance or relevant biological variance. Indeed, ICC values increased overall with PC clocks (Figure 7B) but not to the same extent as with technical replicates (Figures 3-4), reflecting the remaining within-person biological variance due to each person’s longitudinal aging process.

**Figure 7.**
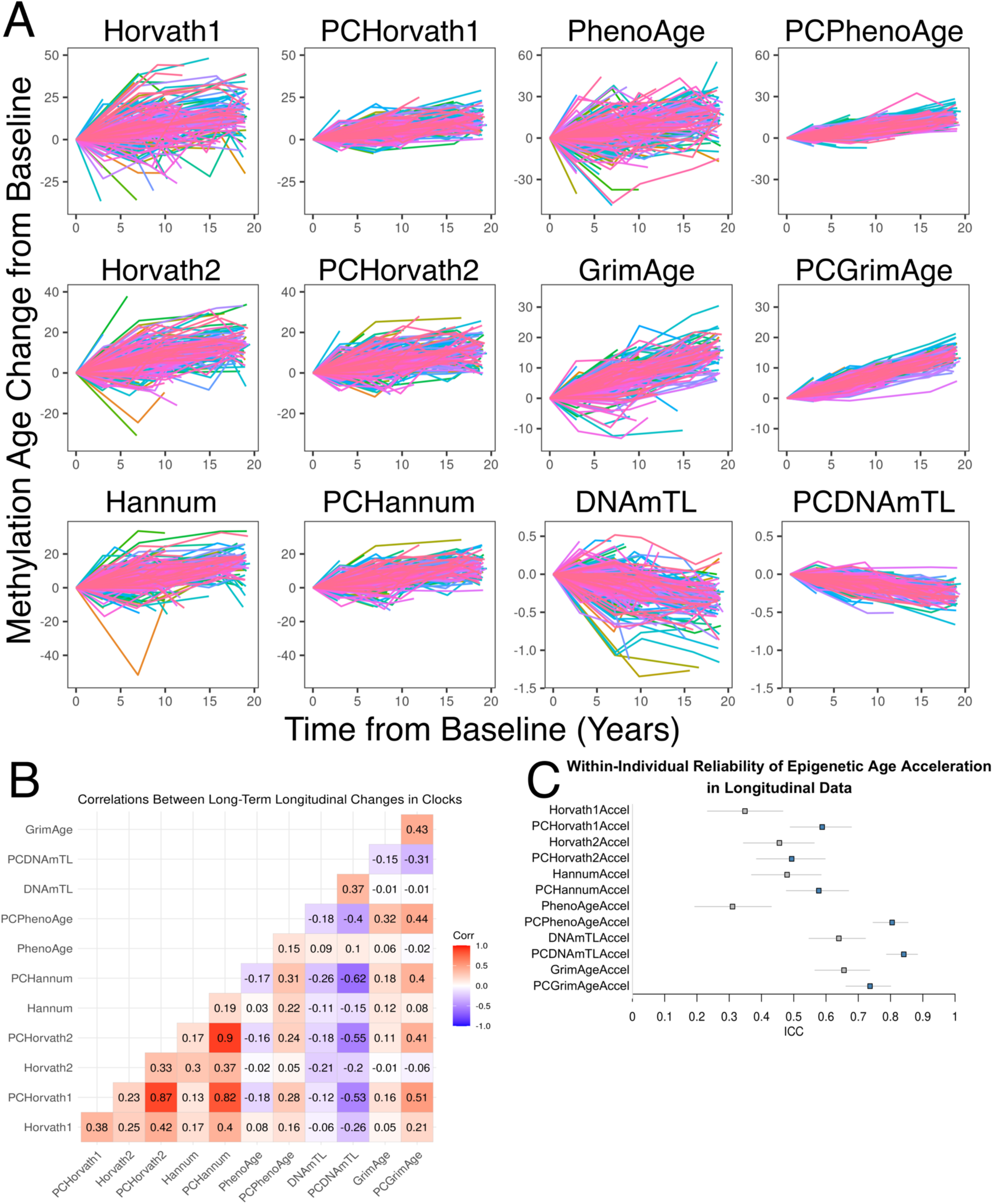
PC clocks show increased stability in long-term longitudinal blood DNAm data. (A) Each line shows the trajectory of an individual’s epigenetic age relative to their baseline during the follow-up period. Colors are included primarily to distinguish between different individuals. (B) Slopes were calculated for every individual for each clock to obtain correlations in short-term longitudinal changes in the clocks. (C) ICC values reflect within-individual variance relative to total variance for each clock.

We also examined short-term longitudinal data with 133 combat-exposed military personnel (baseline age range 18-54) from the Prospective Research in Stress-related Military Operations (PRISMO study) (van der Wal et al. 2020). DNAm measurements were performed at baseline pre-deployment and then at 1 or 2 additional time points post-deployment, up to 500 days follow-up after baseline. The original CpG clocks fluctuated dramatically, deviating up to 10-20 years off the average trajectory (Figure 8A). The PC clocks improved upon this modestly, but there was still significant fluctuation (Figure S19). We hypothesized that this was related to changes in cell composition, which can change as a result of stress, diurnal rhythms, sleep, and other factors (Lasselin et al. 2015; Ackermann et al. 2012). Thus, cell composition could be affected either by combat exposure, or by day-to-day and time-of-day variation. Indeed, most PC clocks showed greatly improved stability after adjusting for cell counts, though this improvement was not seen for the CpG clocks (Figure 8A, Figure S19). Thus, this relevant biological variation only becomes apparent after removing technical noise. Again, longitudinal changes in PC clocks were far more correlated than with the CpG clocks (Figure S20). For example, correlation of longitudinal changes in PhenoAge and GrimAge increased from r = 0.1 to r = 0.85 using PC clocks, while correlation between Horvath1 and Horvath2 increased from r = 0.01 to r = 0.86. ICCs also tended to be higher for the PC clocks and after cell adjustment in most cases, with a few exceptions (Figure S21). Despite increased apparent stability in trajectories, DNAmTL and GrimAge ICCs decreased after cell count adjustment, suggesting some of their primary aging signal is driven by changes in cell counts. For GrimAge, the PC clock version actually had a lower ICC in short-term longitudinal data. The reason is unclear, but it is possible that stress or other changes during the study period drove *bona fide* changes in PCGrimAge for some individuals.

**Figure 8.**
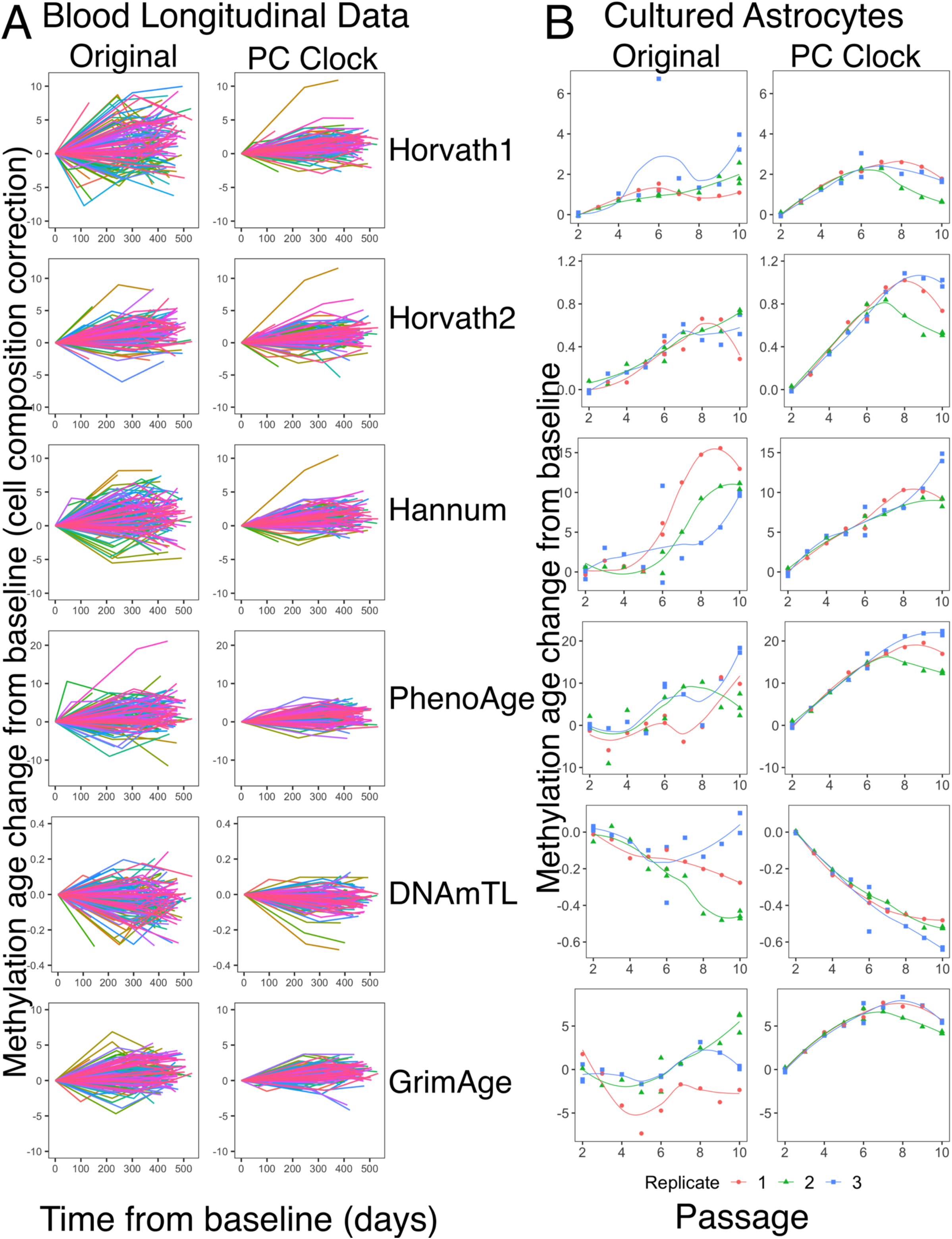
PC clocks show increased stability in short-term DNAm data. (A) Short-term longitudinal blood DNAm data was measured with up to 500 days follow-up. Each line shows the trajectory of an individual’s epigenetic age relative to their baseline during the follow-up period. Cell-adjusted trajectories were adjusted based on proportions of 7 cell types imputed from DNAm data (see methods). (B) DNAm from astrocytes was measured at every passage in cell culture for 3 replicates. Each curve shows the trajectory of one replicate over time from baseline. The baseline is the mean between the replicates at the first DNAm measurement.

We also replicated the increased stability of PC clock trajectories in short-term longitudinal data in a cohort of 13 schizophrenia patients treated with clozapine, measured at 2-3 time points over 1 year including just prior to clozapine initiation (Figure S22). DNAmTL increased during this period (p = 0.0226, 121 bp/year), but PCDNAmTL did not (p = 0.320, 22 bp/year) (Table S13). DNAmTL’s increase was likely due to a combination of noise and small sample size. Thus, the PC clocks may be useful in avoiding false positives in small pilot studies of interventions targeting epigenetic age.

### PC clocks show improved stability in an *in vitro* model of aging

*In vitro* models of aging are useful for experimental investigation of cellular aging and senescence, and for screening novel pharmacological interventions (Itahana et al. 2004; Chen et al. 2013). Some patterns of epigenetic aging are shared by *in vitro* and *in vivo* contexts, suggesting epigenetic clocks can be readouts for aging in cell culture (Liu et al. 2020; Wagner 2019). We derived 3 lines of primary astrocytes from the cerebral cortex of the same fetal donor. These astrocytes were cultured for 10 passages under normoxic conditions (20% O_2_), and measured DNAm at each passage (Figure 8B). While the original CpG-based clocks detected some changes in epigenetic age, there was substantial deviation among replicates and fluctuations between time points. In contrast, the PC clocks showed strong agreement between replicates and smooth increases in epigenetic age up to passage 6. Beyond passage 6, the rate of change slows down (even reversing for some clocks) and the replicates diverge, coinciding with a divergence in the rate of population doubling (Figure S23). This may be biologically significant, reflecting cell death with mortality selection, cellular senescence, or adaptation to the culture environment. Overall, the reliability of PC clocks makes them more useful than the original clocks as biomarkers for *in vitro* aging.

## Discussion

Convenient technology, accessible data, and ease of calculating epigenetic clocks have led to a boom in studies investigating associations between epigenetic age and aging outcomes or risk factors (reviewed in Horvath & Raj 2018; Fransquet et al. 2019; Ryan et al. 2020). More recently, some studies have also begun to apply epigenetic clocks to assess the effects of aging and longevity interventions (Chen et al. 2019; Fahy et al. 2019). However, little attention has been paid to the reliability of these clocks until recently. Our data shows that unreliable probes contribute to significant technical variability of epigenetic clocks. Even if a pair of replicates agree on a given clock, this occurs only because noise from different CpGs cancels out by chance, and that same pair of replicates will usually deviate for other clocks. The magnitude of this unreliability is large, causing up to 3 to 9 years difference between some technical replicates (Figure 1). For comparison, the standard deviation of epigenetic age acceleration is 3 to 5 years depending on the clock. This acceleration value is what predicts aging outcomes such as mortality above and beyond what chronological age can predict. Epigenetic age acceleration is contaminated by technical variation, hampering the utility of epigenetic clocks and potentially leading to both false positive and false negative results.

We present a computational solution to reduce this technical variability, by training clocks using principal components instead of individual CpGs. PCA can extract the shared aging signal across many intercorrelated CpGs while ignoring noise. The resulting PC clocks are highly reliable even though they do not require *a priori* knowledge of CpG ICCs for their construction (Figures 4-5). This is particularly important because replicate data does not exist for many cohorts, tissues, and aging phenotypes. This method can even increase the reliability of clocks that already have high ICCs (e.g. GrimAge). At the same time, the PC clocks preserve the relevant aging signals unique to each of their CpG counterparts (Figure 6); therefore, they reduce technical variance but maintain relevant biological variance. PCA is a commonly used tool and does not require specialized knowledge, and thus this approach is accessible and readily adaptable to improving the reliability of any existing or future epigenetic biomarker.

It is possible that future technology or processing methods may improve the reliability of individual CpGs, but the question remains what to do about epigenetic clock reliability now, given the massive amount of data collected on methylation arrays by current aging cohort studies. Using reliability information during training modestly improves reliability for CpG-based clocks (Figure 1K) but does not add anything to PC-based clocks (Table S11). This suggests that the PC-based clocks may be near the possible limit of reliability, at least using linear techniques. It will be interesting to see if more advanced non-linear machine learning methods can further improve reliability, though these would be less accessible and more complex to use.

PC clock reliability will be critical for employing the epigenetic clocks for personalized medicine and clinical trials. For example, it would be highly misleading for a conventional CpG-based epigenetic clock to indicate that a person has aged 9 years over the course of 1 day, if the difference is solely attributable to technical variation. Measurement changes in biological age after starting a new treatment or lifestyle change would not be trustworthy. While the residual noise in PC clocks (1-1.5 years maximum) may still pose challenges for detecting small effect sizes, these may be overcome by studies of sufficient sample size or those with intervals longer than 1 year.

Fluctuations in cell counts contribute to clock variation in short-term longitudinal studies, so these studies should account for any factors that affect cell composition, such as stress, sleep, or time of day. However, these efforts would be hindered if technical variation is not first addressed (Figure 8).

Our study has implications for aging biology in general. First, CpGs (or other biological variables) with low reliability show reduced associations with aging phenotypes, but this is a technical artifact where the signal is contaminated by noise. This presents a systematic bias in the aging literature where some CpGs may be ignored simply because they are harder to measure, even though they are biologically relevant. These can be thought of as false negative results. This issue may be mitigated by focusing on concerted changes across many CpGs, rather than studying one at a time. PC clocks utilize CpGs across the ICC spectrum (Figure S17), suggesting that PCA can extract relevant information from low-ICC CpGs while ignoring noise. Second, the specific identities of CpG sites (and associated genes) included the epigenetic clock may be less important than previously supposed. After all, elastic net regression selects only a small subset of CpGs for traditional clocks from a larger group of multicollinear CpGs. Instead, it may be better to conceptualize the epigenetic clock as measuring global processes affecting many CpGs and genes in concert, reflected in the covariance captured by PCA. Aging involves numerous intercorrelated changes in many bodily systems over time that leads to dysfunction, and therefore focusing on a small set of specific variables may miss the forest for the trees.

Overall, we were able to drastically improve the technical reliability of epigenetic clocks, while simultaneously maintaining or even increasing their validity. Moving forward, these measures may provide critical tools for assessing aging interventions, tracking longitudinal aging trajectories, and gleaning biological insights from global shifts in DNAm patterns over the lifespan.

## Methods

### Reliability analyses and datasets

We calculated ICC using the icc function in the irr R package version 0.84.1, using a single-rater, absolute-agreement, two-way random effects model, after consulting guidelines from Koo and Li (Koo & Li 2016). Two-way was chosen because all subjects were measured by the same raters (e.g. two batches of replicates). The random effects model allows reliability results to generalize to other DNAm batches. Absolute agreement was used because we aim not only for methylation age to correlate between batches but also for their values to agree. Single rater was used because usually methylation age is based on a single sample rather than the mean of multiple samples. ICCs less than 0 were sometimes re-coded as 0, either for figure presentation purposes or to compare the ICCs to previous datasets where this re-coding was done.

To assess CpG and clock reliability, we used the publicly available dataset GSE55763 from Lehne and colleagues (Lehne et al. 2015), consisting of 36 whole blood samples measured in duplicate from the London Life Sciences Prospective Population (LOLIPOP) study. We selected this sample because it had the widest age range (37.3 to 74.6 years) of available replicate datasets, which is important for assessing epigenetic clock performance. The sample size was sufficient, as Sugden et al. found that running just 25 pairs of replicates was sufficient to identify 80% of reliable probes (Sugden et al. 2020), and our individual CpG ICCs were broadly in agreement with a larger sample of 130 sets of replicates with a narrower age range (Bose et al. 2014). The replicates in GSE55763 were done in separate batches to maximize the impact of technical factors. The dataset had been processed using quantile normalization which Lehne et al. found showed the best agreement between technical replicates out of 10 normalization methods. It was also adjusted using control probes to remove systematic technical bias (e.g. from batches and plates). Note that none of the 78,464 CpGs we analyzed in-depth had any missing values in GSE55763.

### Epigenetic Biomarkers

Epigenetic biomarkers were calculated either according to published methods, or using the online calculator (https://dnamage.genetics.ucla.edu/new) to obtain GrimAge and cell proportions. For nomenclature, we referred to clocks that predict chronological age by the last name of the first author of the publication that first reported them. For those that predicted other phenotypes, we used the descriptive name provided by the original publication. For Figure 7, we adjusted for cell composition using a linear model regressing epigenetic age on chronological age, plasmablasts, CD4+ T cells, exhausted CD8+ T cells, naïve CD8+ T cells, natural killer cells, granulocytes, and monocytes, as previously described (Horvath & Levine 2015).

### Analyses

Analyses were performed in R 4.0.2 and RStudio 1.3.1093. Figures were made using R packages ggplot2 v3.3.3, forestplot v1.10.1, ggcorrplot v0.1.3, pheatmap v1.0.12, or WGCNA v1.70-3. Correlations were calculated using biweight midcorrelation from the WGCNA package, unless otherwise stated. Mortality in FHS was calculated using the survival v3.2-7 package.

### Training proxy PC clocks

We trained principal components (PCs) in different datasets for each clock (Table S7). Each dataset (beta matrices) was filtered down to 78,464 CpGs that were (1) on the 450K array, EPIC array, and a custom array (Elysium), and (2) shared by our datasets used for PCA training, PC clock training datasets, and reliability analysis. We then performed mean imputation for missing values, as imputation method did not appreciably affect PCs due to the large number of CpGs they incorporate. PCA was done using the prcomp function in R with centering but without scaling beta values. Elastic net regression to predict age or CpG clock value from the PC scores were done using the glmnet v4.1-1 package. Alpha value was 0.5, as alpha did not appreciably affect reliability or prediction accuracy. The lambda value with minimum mean-squared error was selected using 10-fold cross-validation. The final PC reported by prcomp was excluded from elastic net regression because it is not meaningful when number of variables is greater than number of samples. Test data were then projected onto the PCs, using the centering from the original training data, allowing for prediction of the outcome variable.

We found we could predict either the original outcome variable (e.g. chronological age) or the original CpG clock value. We decided to create PC clock proxies for Horvath1, Horvath2, Hannum, DNAmTL, and GrimAge. This was particularly important in the case of Horvath1 and Horvath2, where we substituted some datasets (Table S7) as not all of the original data was available. For example, some of the original Horvath1 data was obtained on the 27K array, while our PCs utilized 450K or EPIC data. A few samples were eliminated that showed dramatic discordance between the original Horvath1 and Horvath2 value and annotated age.

While the predicted ages and age acceleration values for PC clocks and their corresponding CpG clocks always correlated strongly, we found the intercept was sometimes different, leading to a systematic offset in some datasets. However, the CpG clocks themselves often have highly variable intercepts between datasets, which seem to reflect batch effects (McEwen et al. 2018; Liu et al. 2020). Since intercepts are not as interesting for aging studies, compared to slope and age acceleration values, it is not a problem that they do not agree. In fact, the PC clocks often reported more reasonable intercepts than the original CpG clocks (i.e. closer to actual chronological age).

To calculate the contribution of each CpG to the final PC clocks, we multiplied the CpG loadings for each PC by the PC weight in the clock, calculated the sum for each CpG, and divided by CpG standard deviation from the PC clock training data.

