## Supplemental for "A computational solution for bolstering reliability of epigenetic clocks: Implications for clinical trials and longitudinal tracking"

**Figure S1. Histograms of intraclass correlation coefficient (ICC) of clock CpGs.**

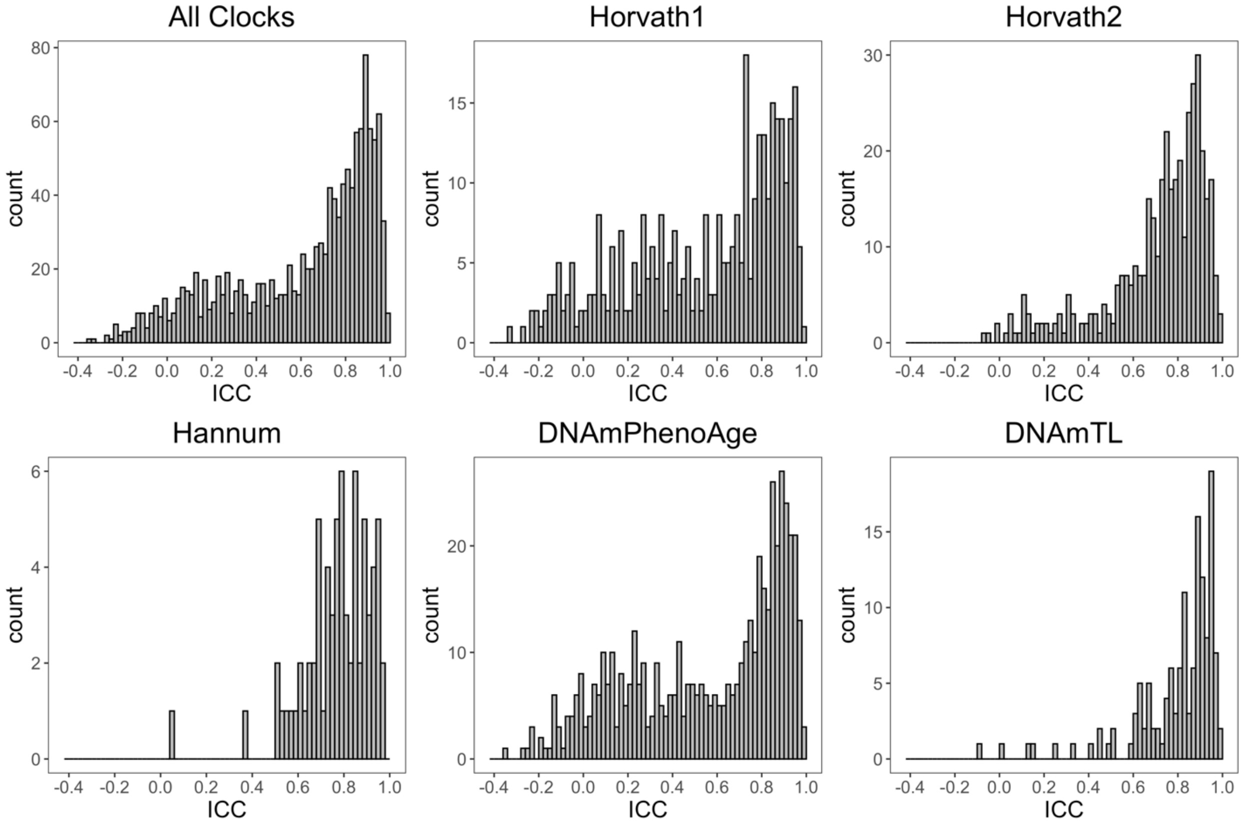


**Figure S2. ICC and agreement of technical replicates for CpG β-values from each clock.** Each point represents one pair of replicates.

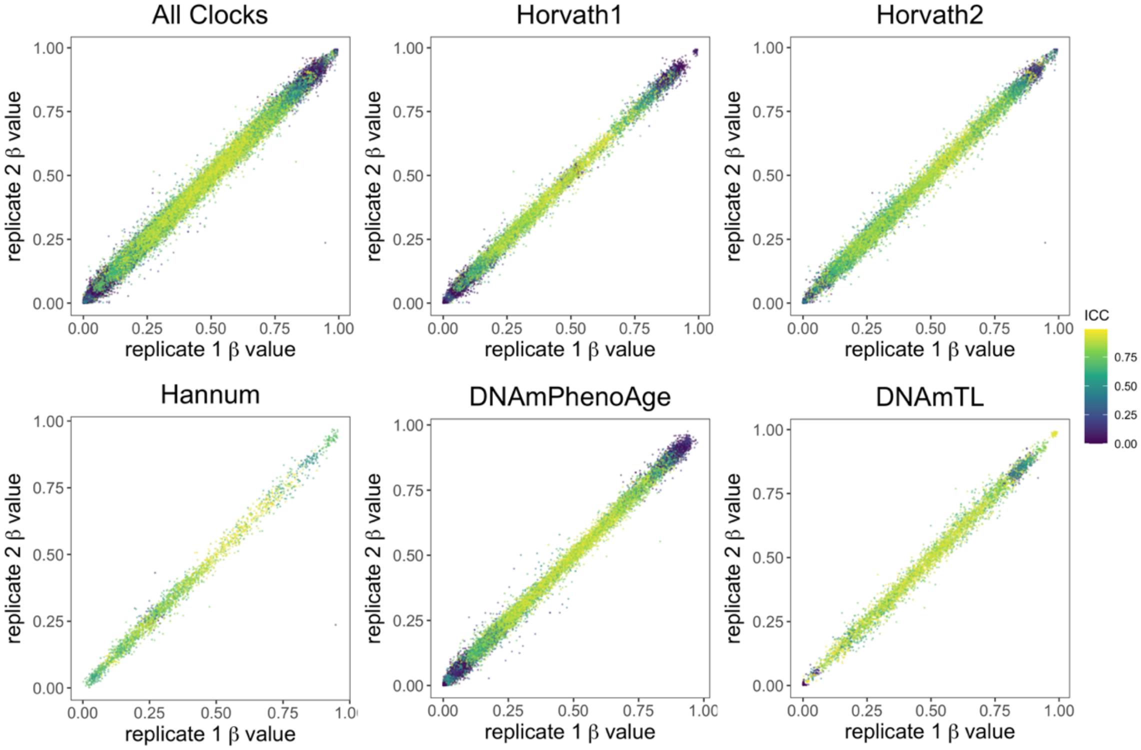

**Figure S3. ICC, mean and standard deviation of CpG β-values in each clock.** Each point is one CpG quantified across 36 samples with 2 technical replicates each.

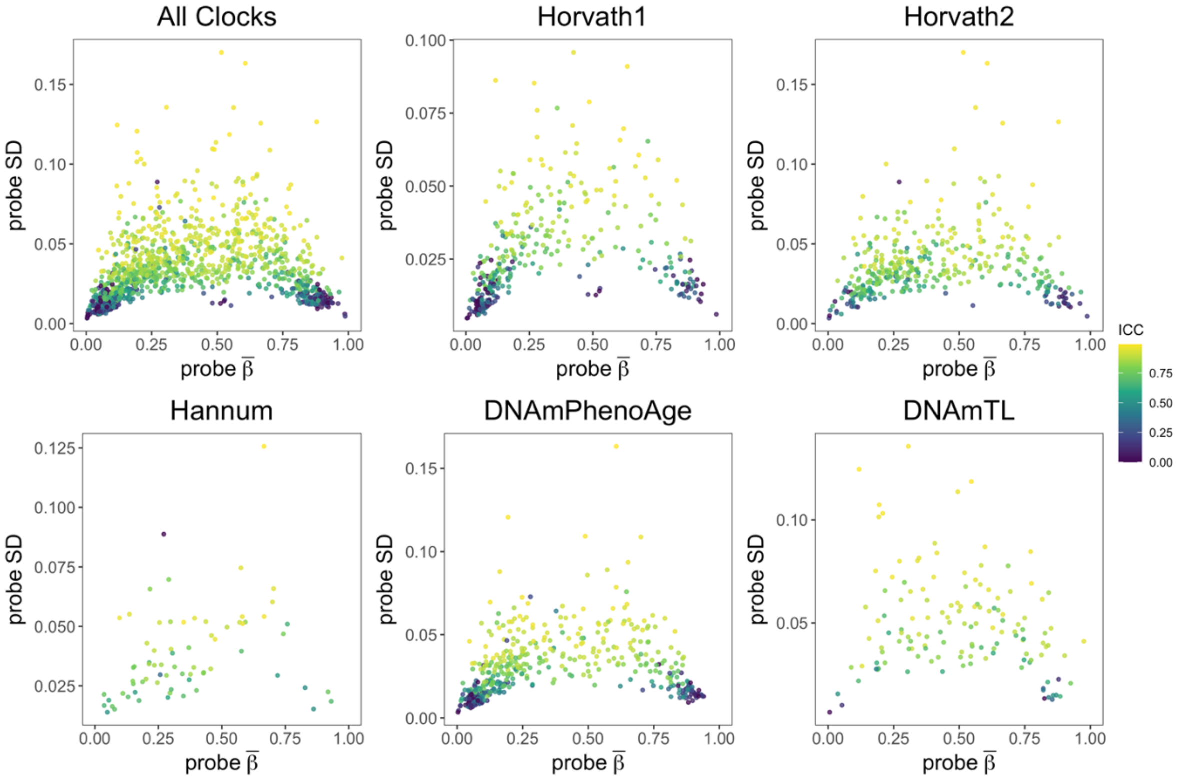


**Figure S4. ICC and standard deviation of CpG β-values in each clock.**

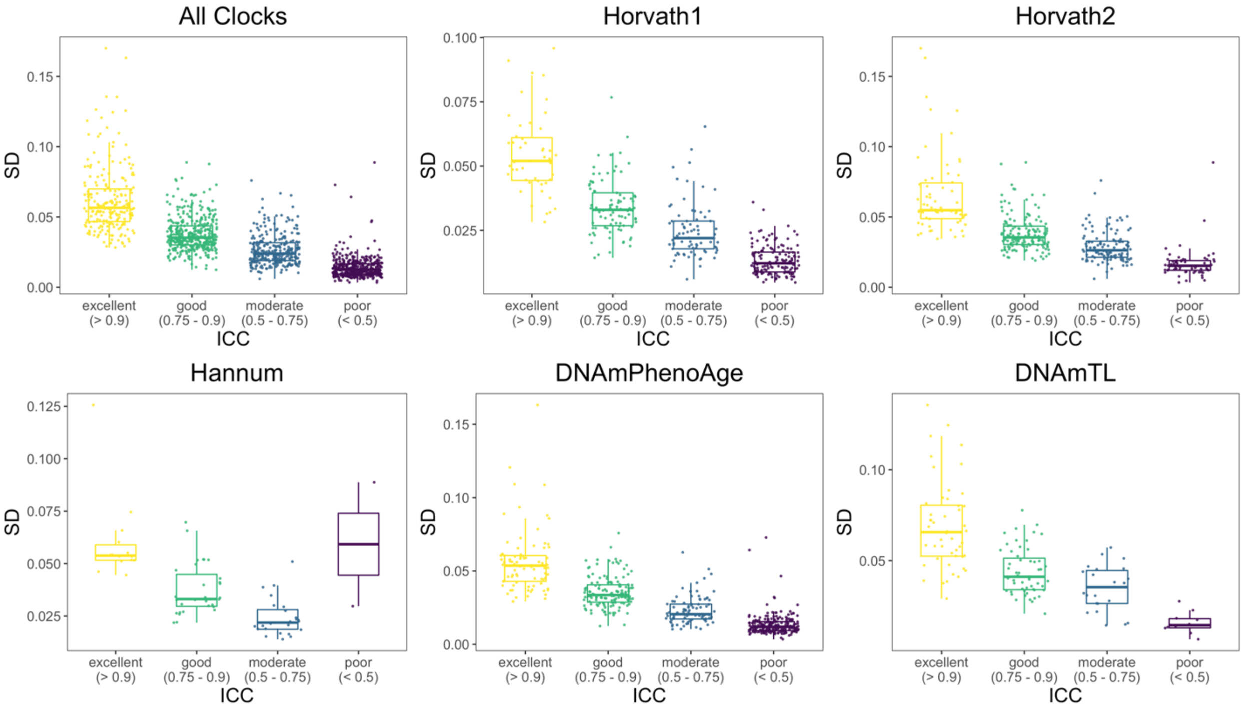

**Figure S5. ICC and mean of CpG β-values from each clock.**

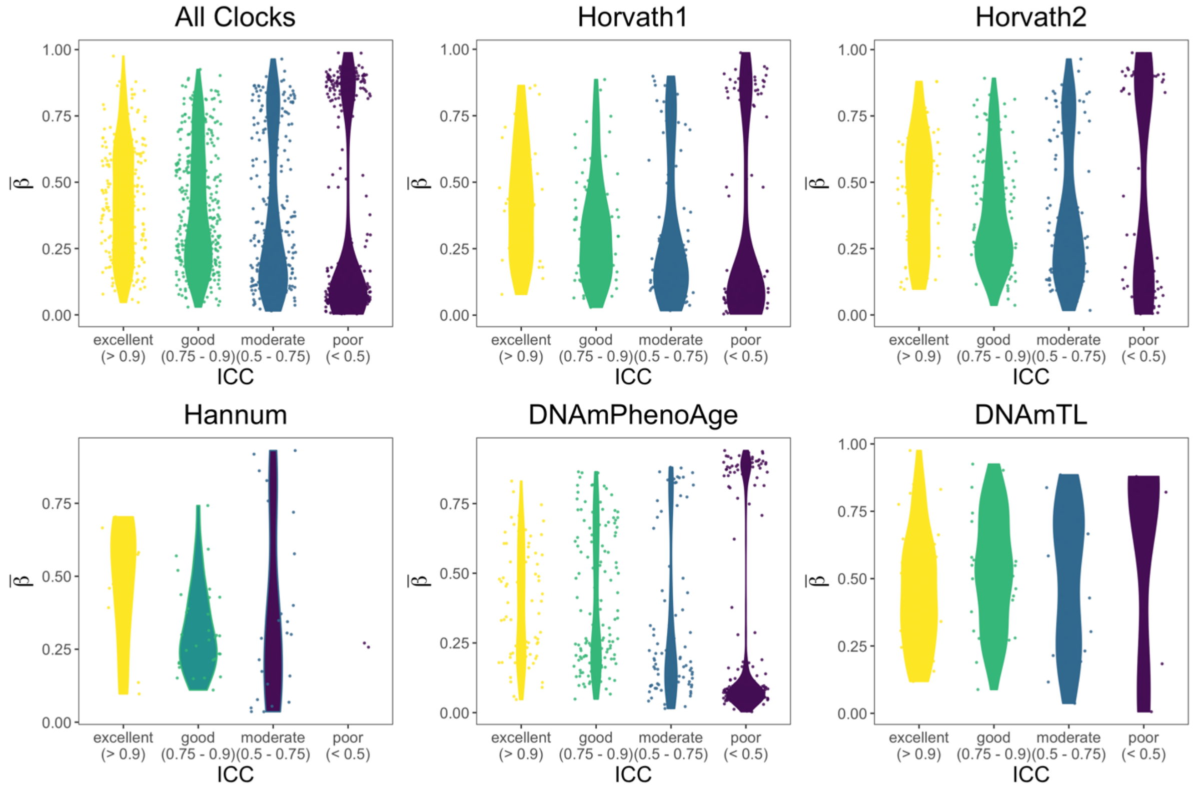


**Figure S6. Age and mortality associations and ICC of CpG β-values from each clock.** Blood age correlations were calculated in GSE40279 (Hannum et al. 2013). Mortality associations (hazard ratios for 1 SD change in β) were calculated in FHS.

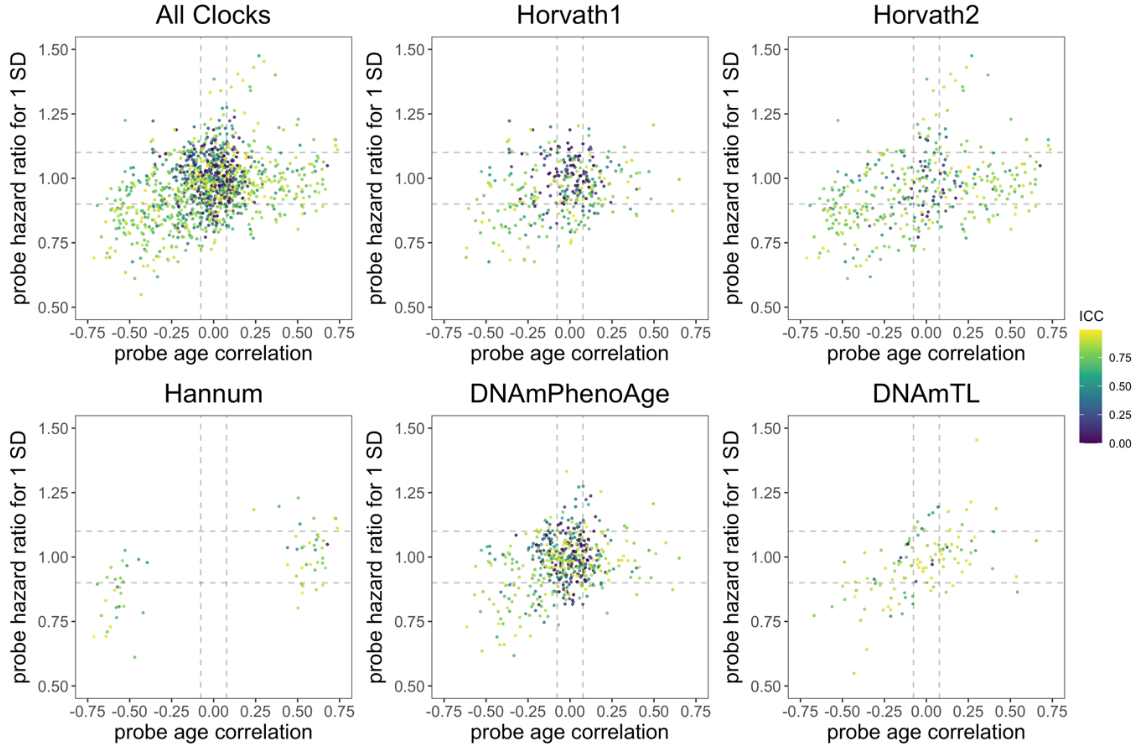

### Figure S7. Age correlation, SD, and ICC of CpG β-values from each clock.

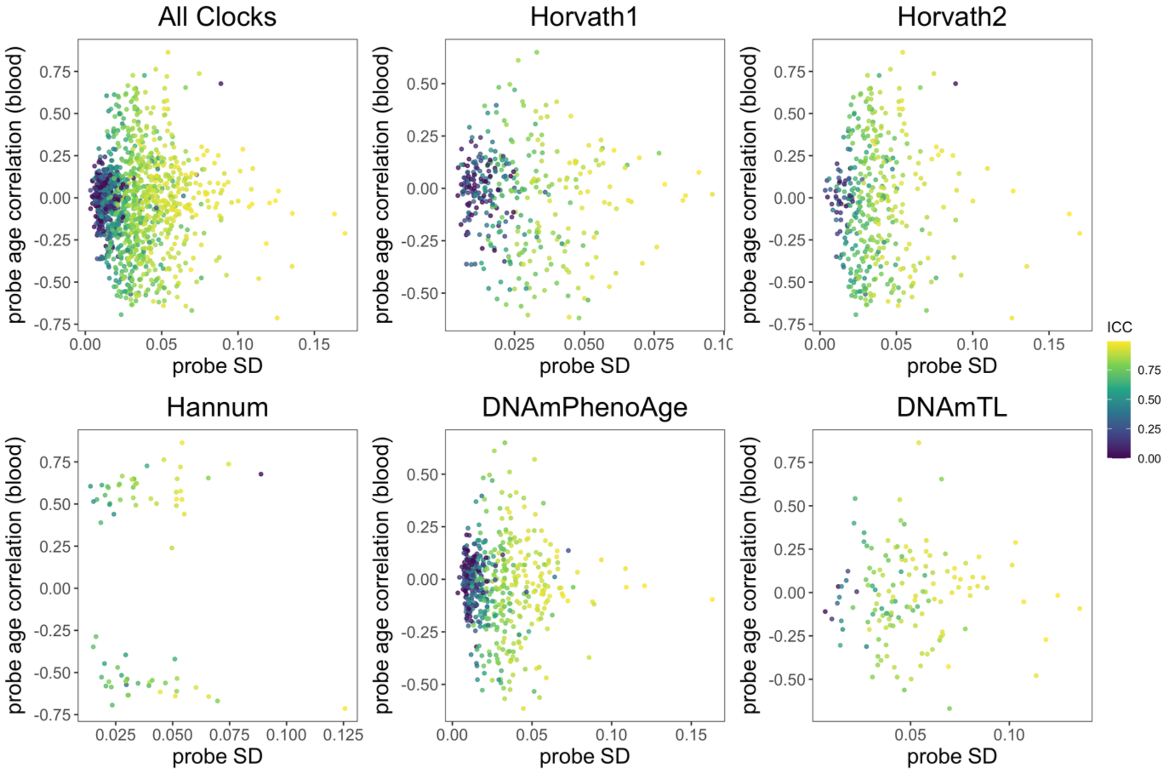


### Figure S8. Age correlation, mean and ICC of CpG β-values from each clock.

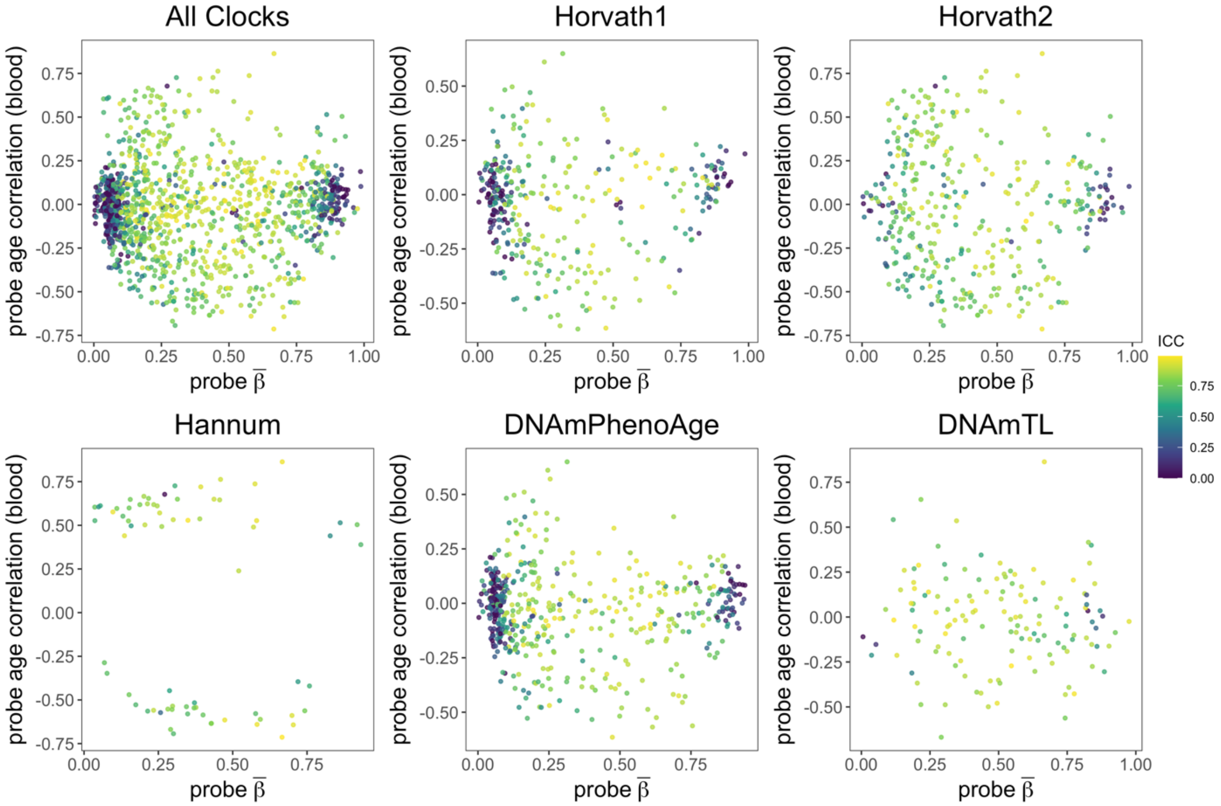

**Figure S9. Agreement of clock values between technical replicates for each clock.** Center black dashed line indicates perfect agreement; gray dashed lines indicate agreement within 2 years. Note that for DNAmTL, a proxy of telomere length, 2 years is equivalent to 0.05 kilobases (kb), as telomere length decreases by 0.025 kb per year.

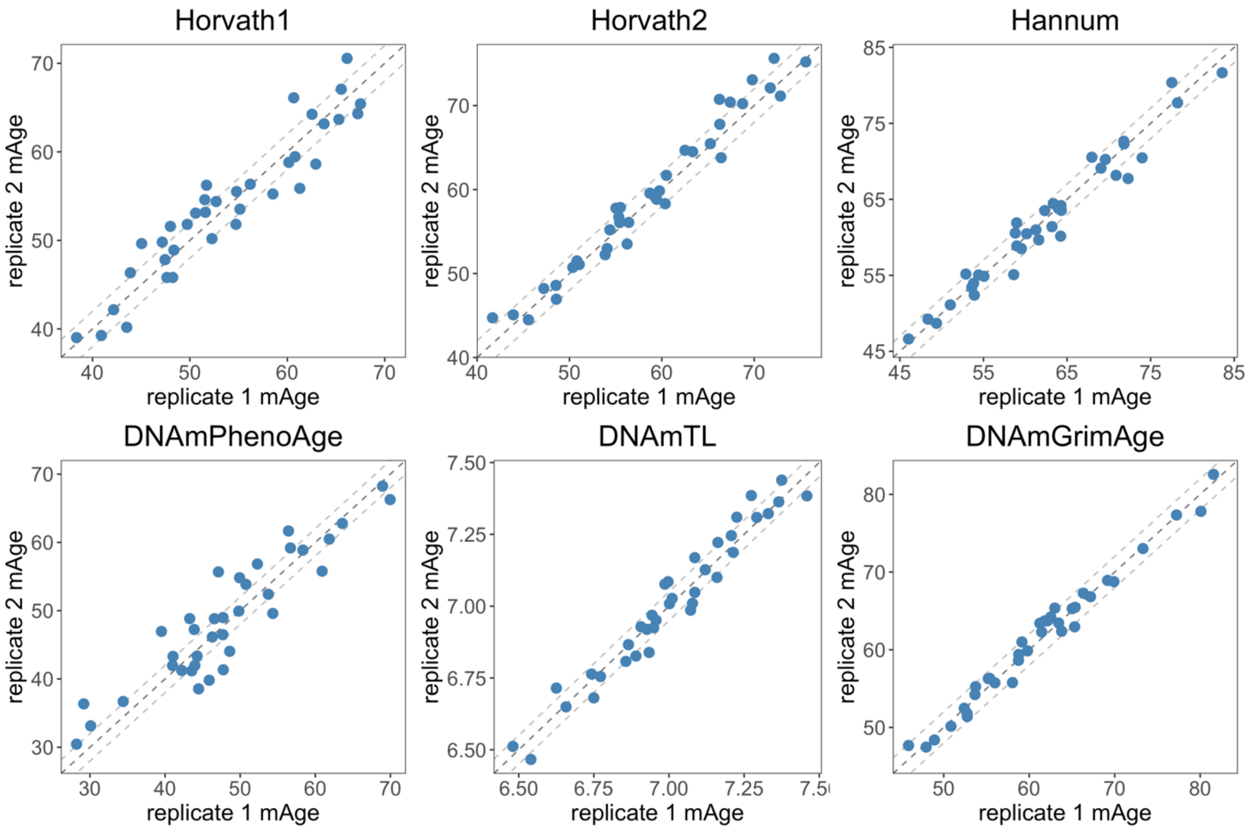

**Figure S10. Histogram of absolute differences between technical replicates for each clock.** We rescaled the axis for DNAmTL, a proxy of telomere length, to reflect that telomere length decreases by ~0.025 kb per year.

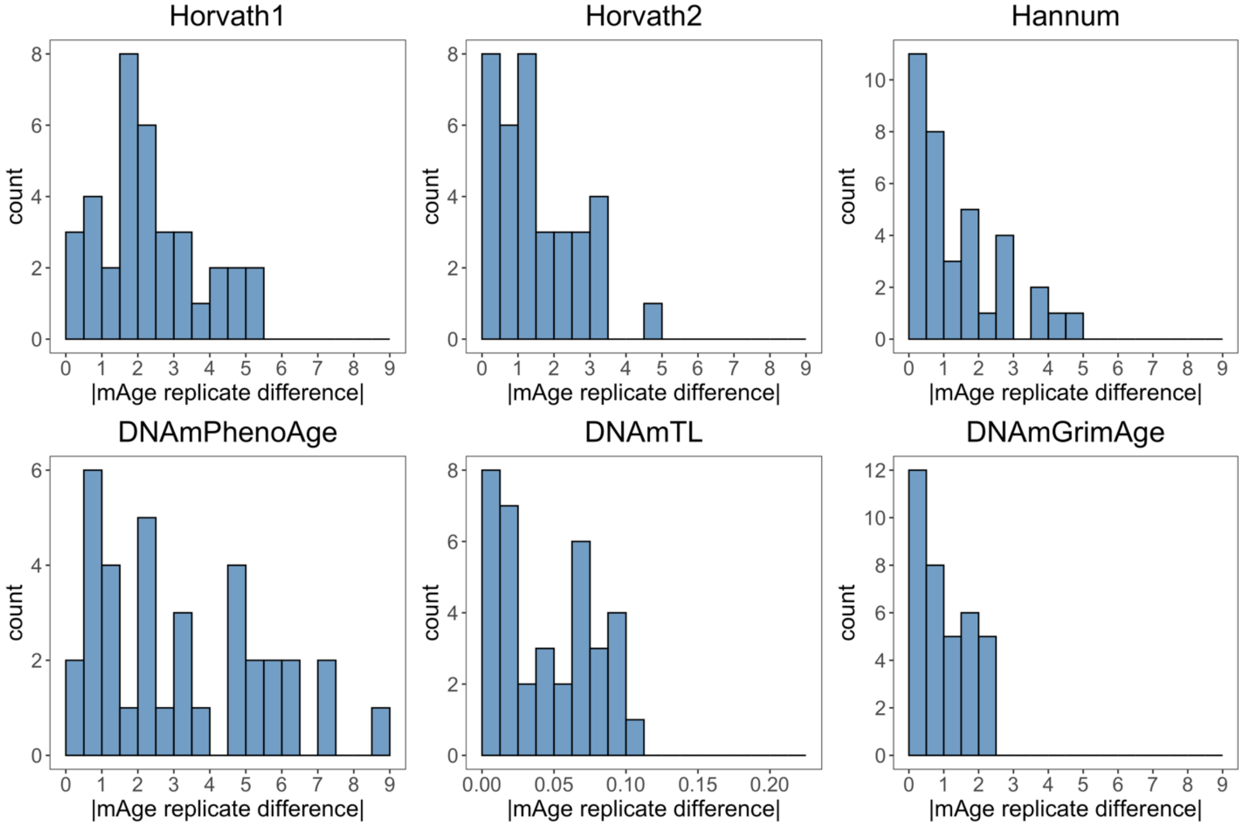

**Figure S11. Correlation plot for epigenetic age differences between replicates.** Epigenetic age replicate differences were calculated for each separately, then the differences were correlated with each other and with age and sex.

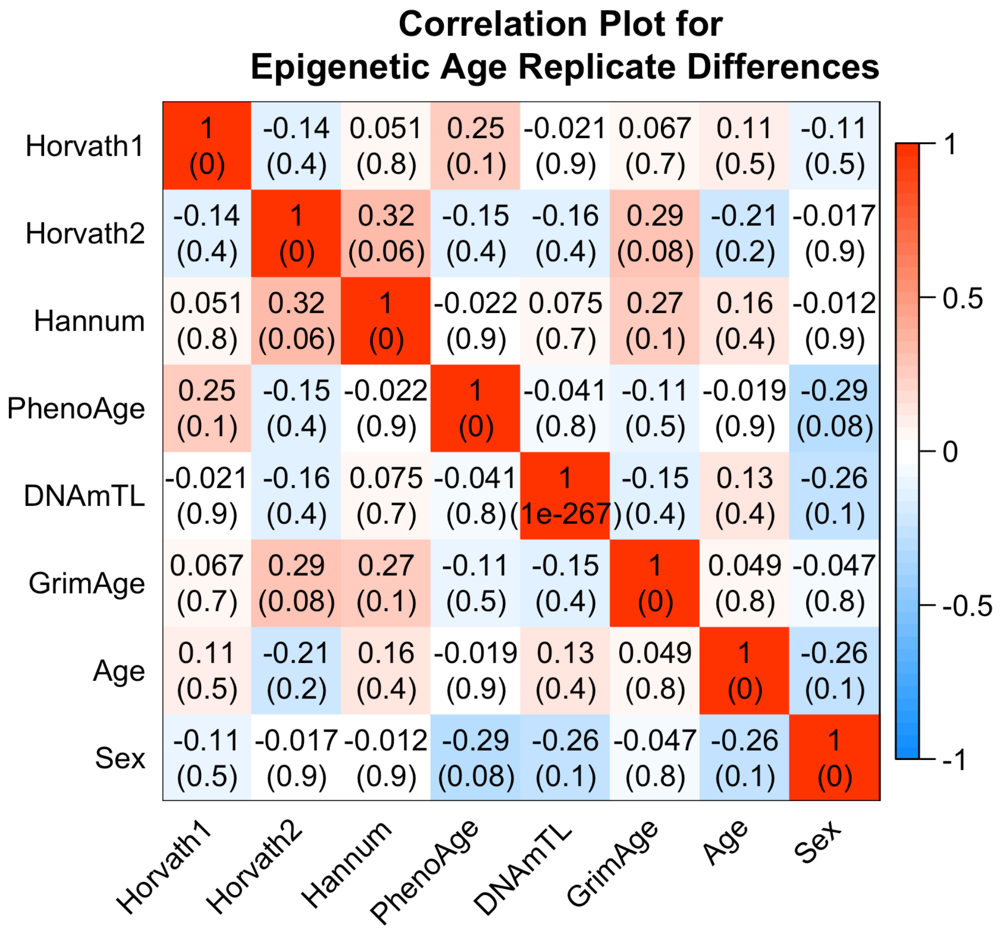

**Figure S12. Heatmap depicting how CpG deviations between replicates contribute to overall epigenetic age scores.** CpG deviation was multiplied by CpG weight within each clock. Rows are CpGs and columns are samples. DNAmTL is in units of base pairs.

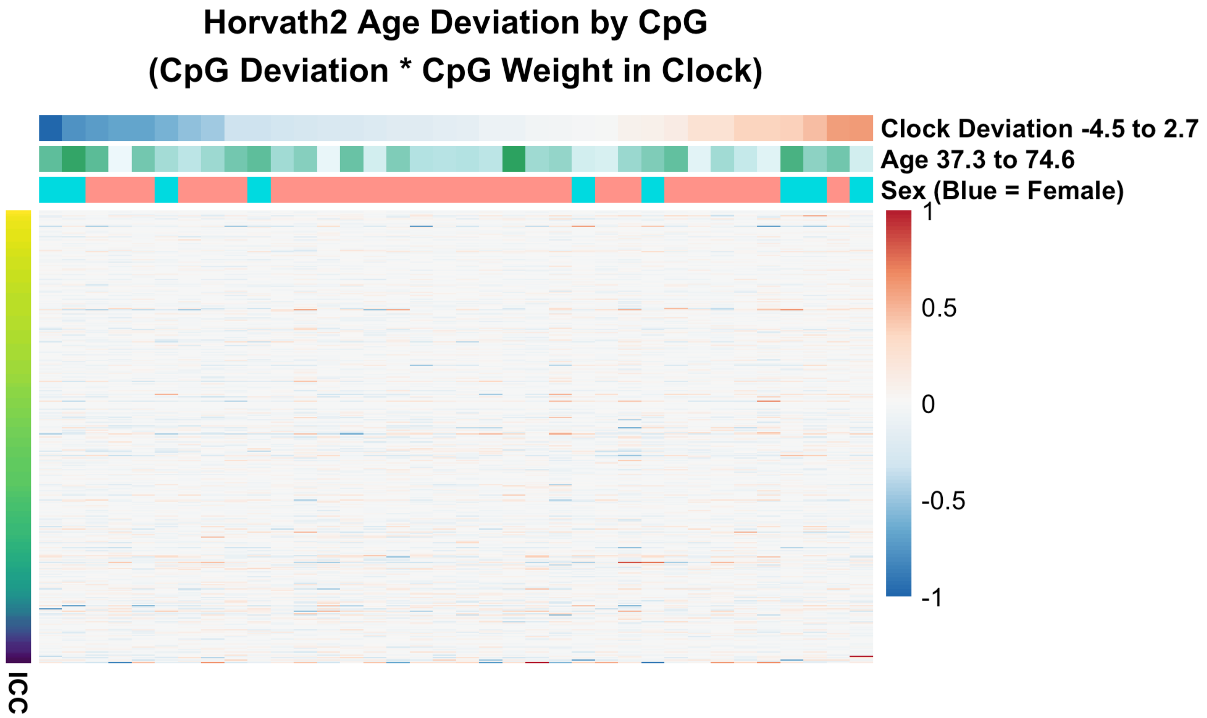


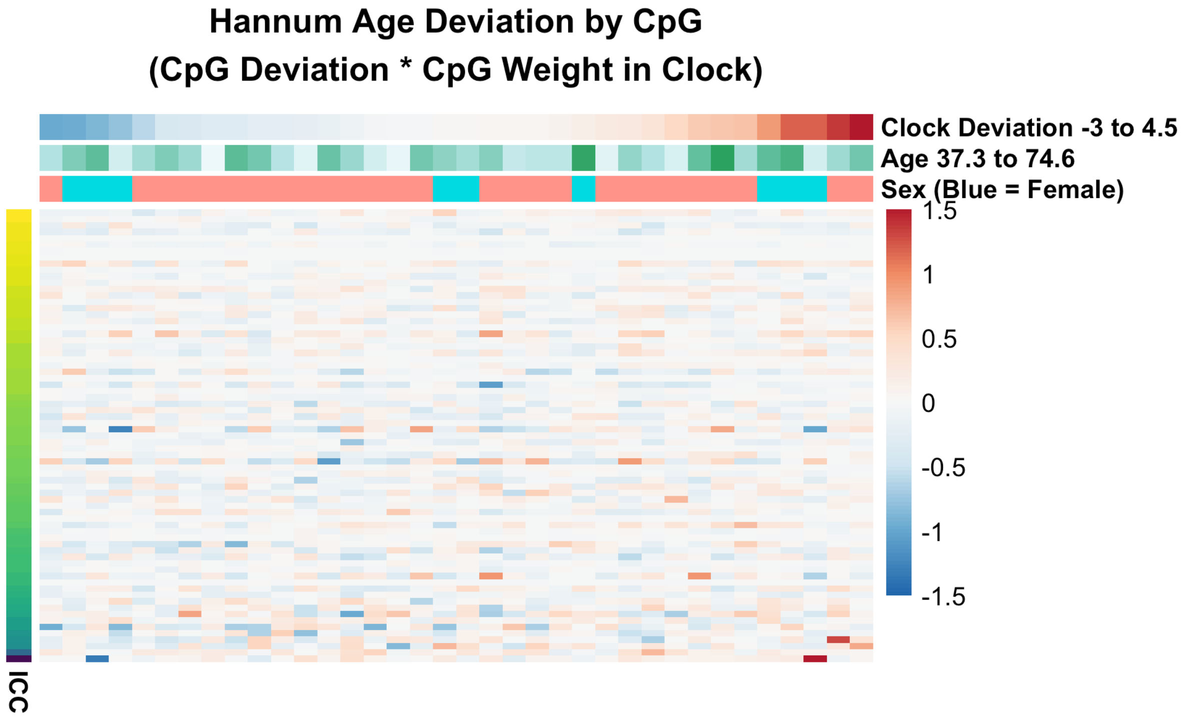

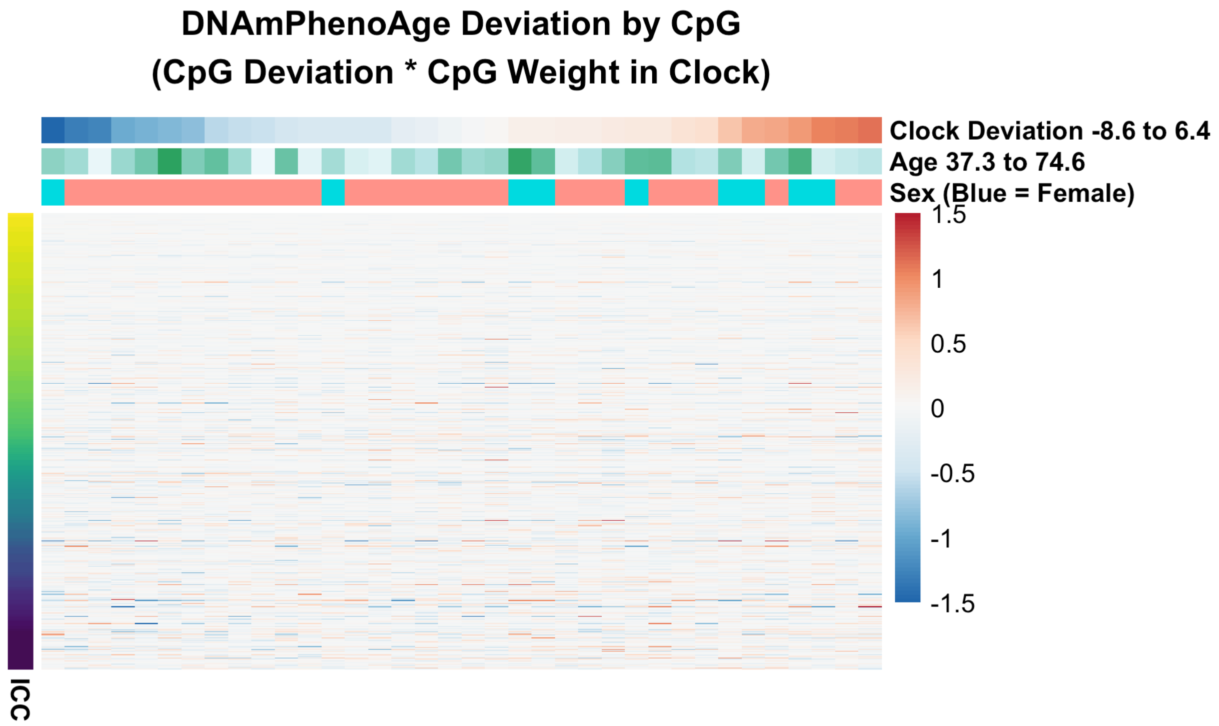
DNAmPhenoAge Deviation by CpG
(CpG Deviation * CpG Weight in Clock)
Clock Deviation -8.6 to 6.4
Age 37.3 to 74.6
Sex (Blue = Female)
1.5
1
0.5
0
-0.5
-1
-1.5
ICC

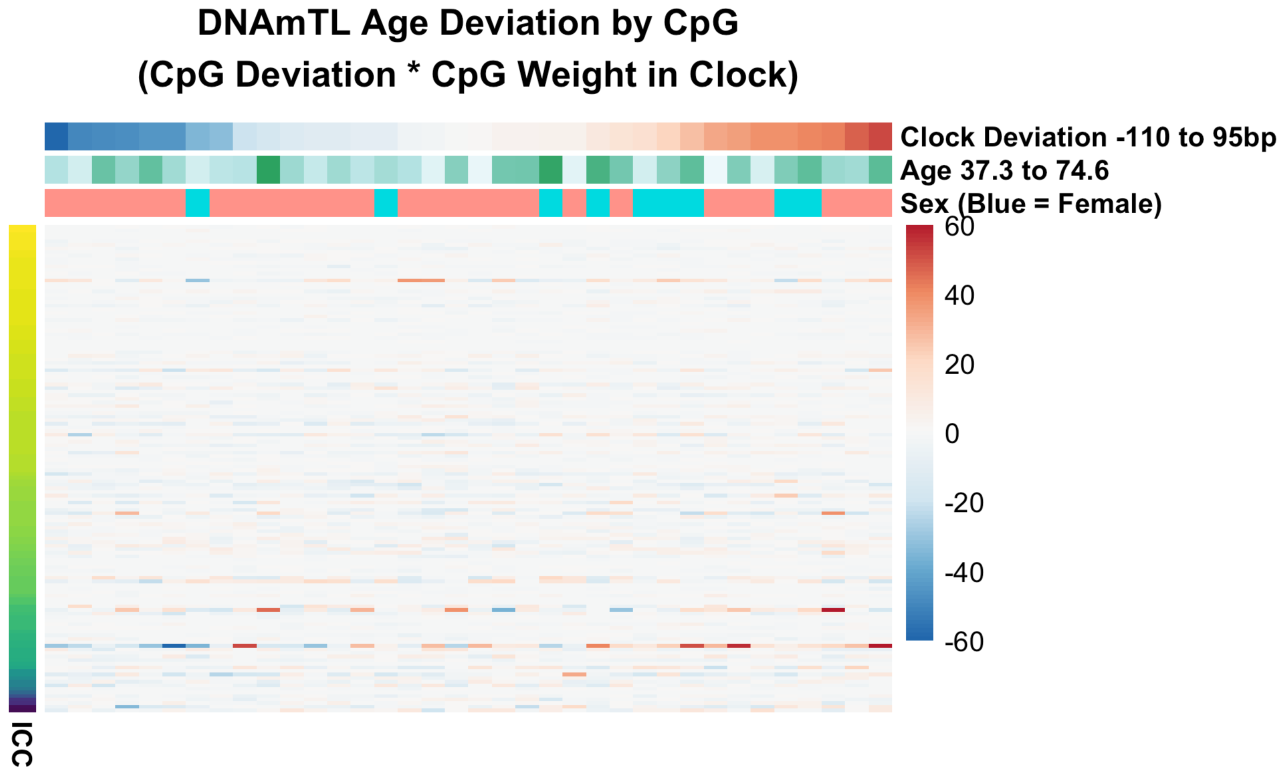
DNAmTL Age Deviation by CpG
(CpG Deviation * CpG Weight in Clock)
Clock Deviation -110 to 95bp
Age 37.3 to 74.6
Sex (Blue = Female)
60
40
20
0
-20
-40
-60
ICC

**Figure S13. Comparison of ICCs for selected 78,464 CpGs to previously published ICC values.** Lehne 2015 is the dataset we analyzed, obtained on the 450K array, age range 37.3-74.6 (Lehne et al. 2015). Bose 2014 was also obtained on the 450K array, age range 45-64 (Bose et al. 2014). Sugden 2020 compared 450K and EPIC, with a cohort of twins where all individuals were 18 years old. Logue 2018 published ICCs for technical replicates on the EPIC array from 11 individuals, drawn from a larger population with mean age 31.8 and SD 8.4. Since Bose 2014 published ICCs with floor value of 0, we changed all Lehne 2015 ICC values less than 0 to a value of 0 to make comparisons consistent. Sugden 2020 or Logue 2018 published negative ICC values, so we adjusted the floor to -0.3 for presentation purposes, though all raw ICC values are in Table S3. The lower ICC correlation with Sugden and Logue may be a combination of differences between EPIC and 450K, and factors that reduce biological variance (e.g. twin design and age range for Sugden; small sample size in Logue).

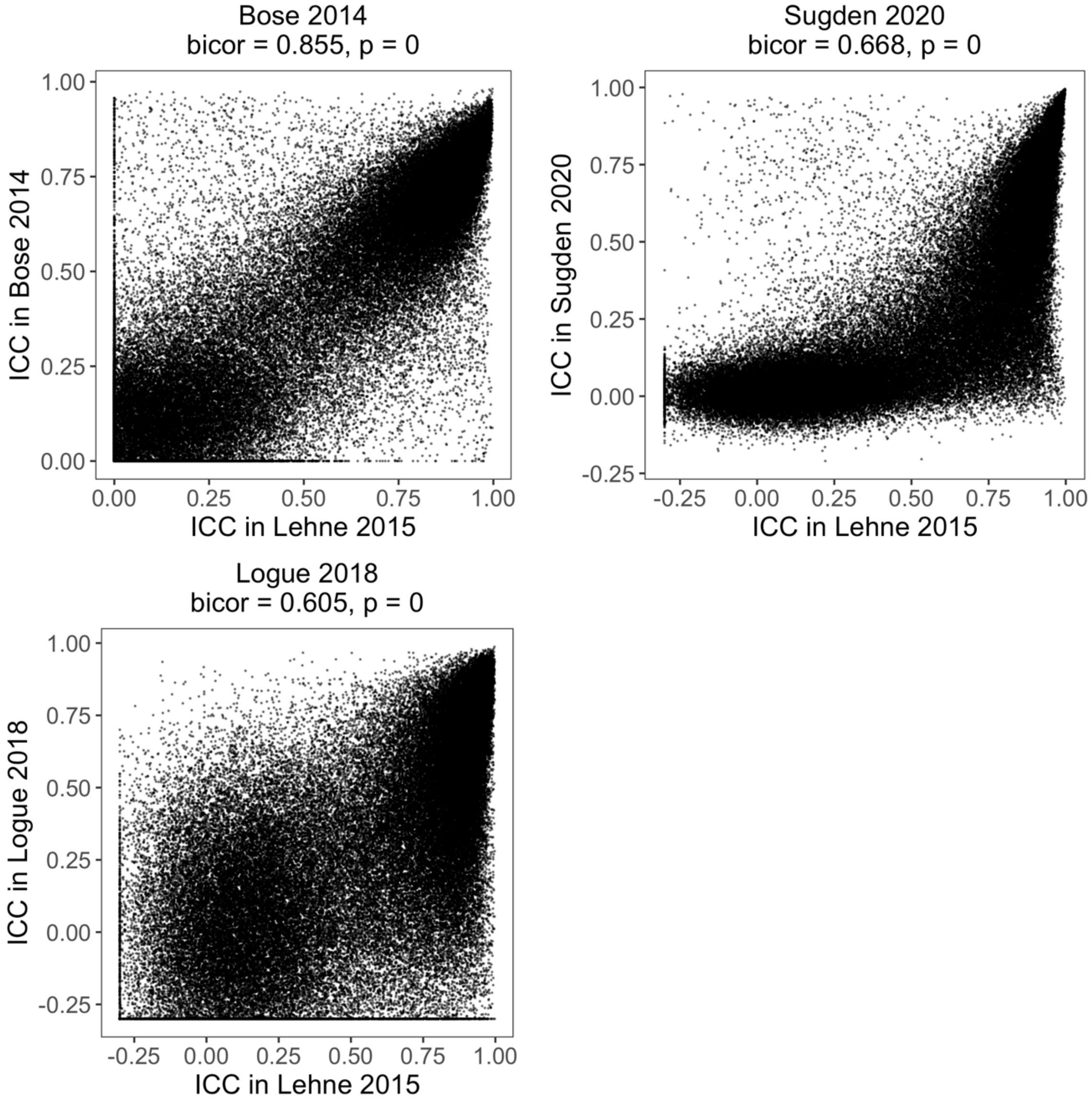

**Figure S14. Age and mortality correlations for CpG ICCs for selected 78,464 CpGs**. Age correlation calculated in GSE40279, and mortality hazard ratio calculated in the Framingham Heart Study after adjusting for age and sex.

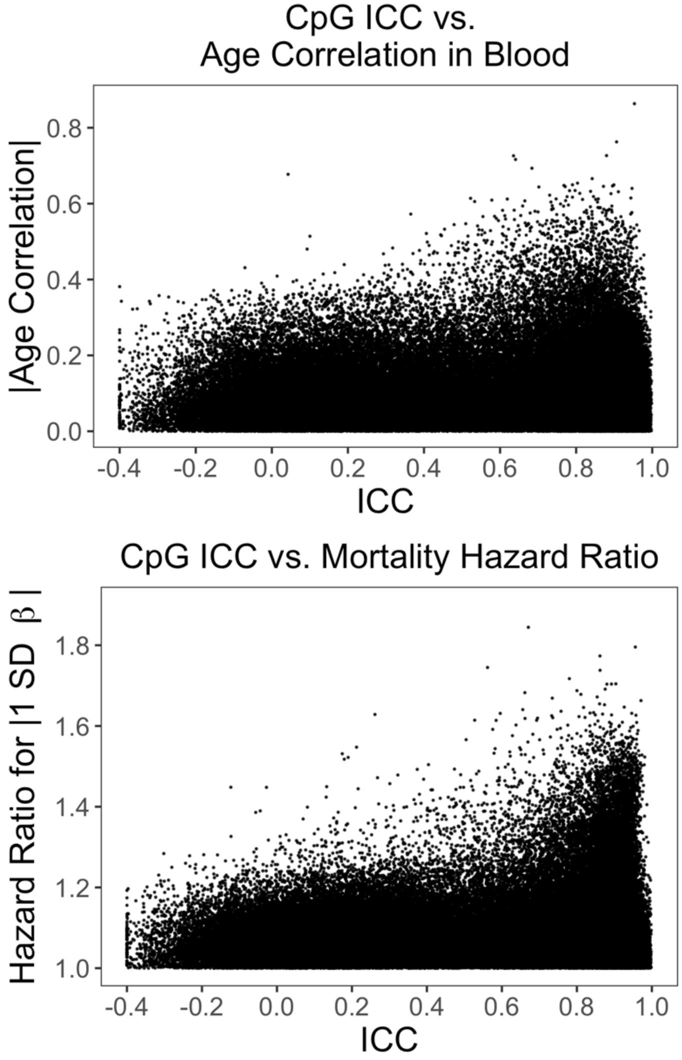

**Figure S15. Analysis of the effect of CpG filtering by ICC on PhenoAge prediction in InCHIANTI.** From the same analysis presented in Figure 2E-F.

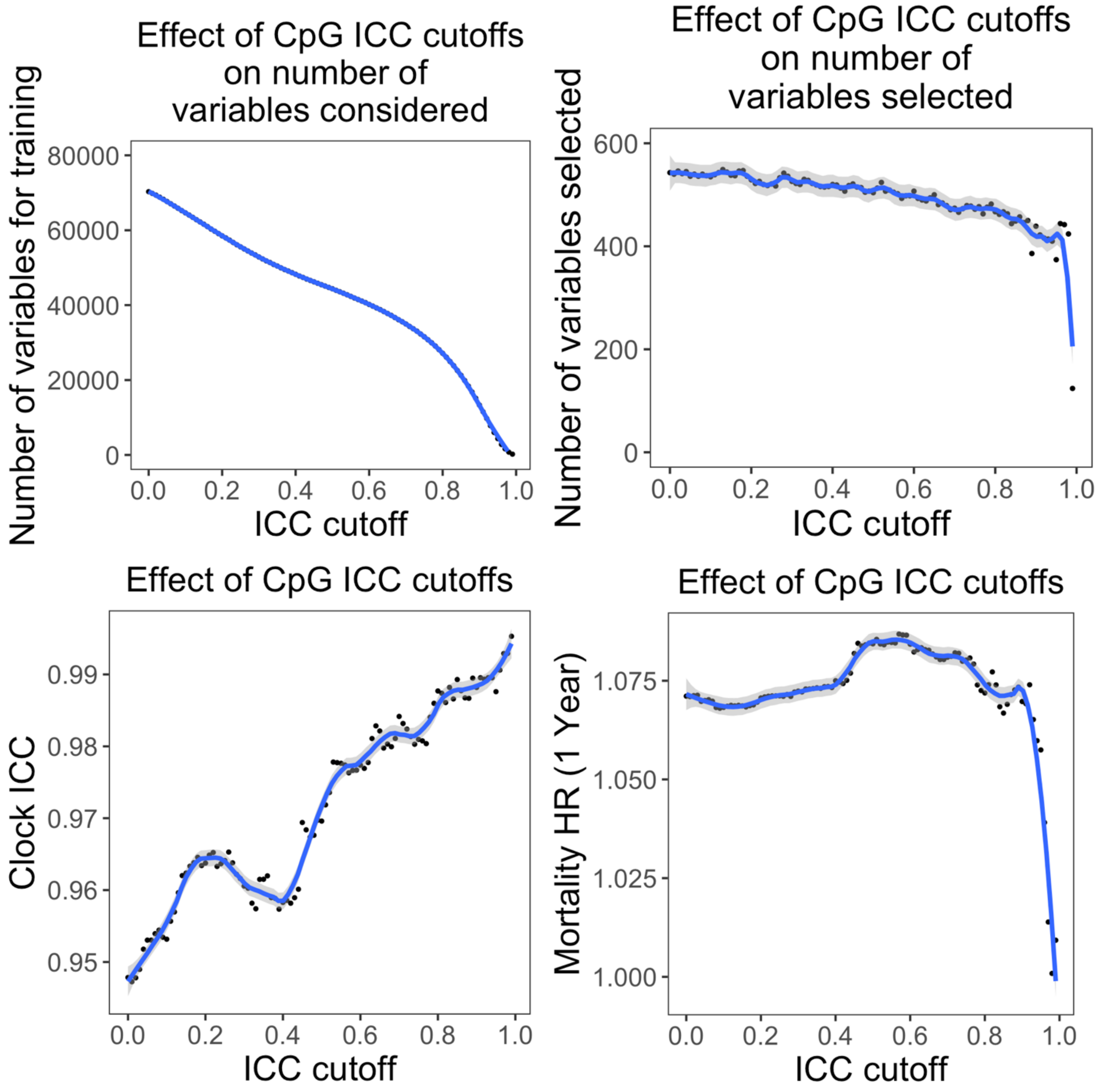

**Figure S16. Analysis of the random CpG subset selection on PhenoAge prediction in InCHIANTI.** This serves as a comparison to filtering by ICC.

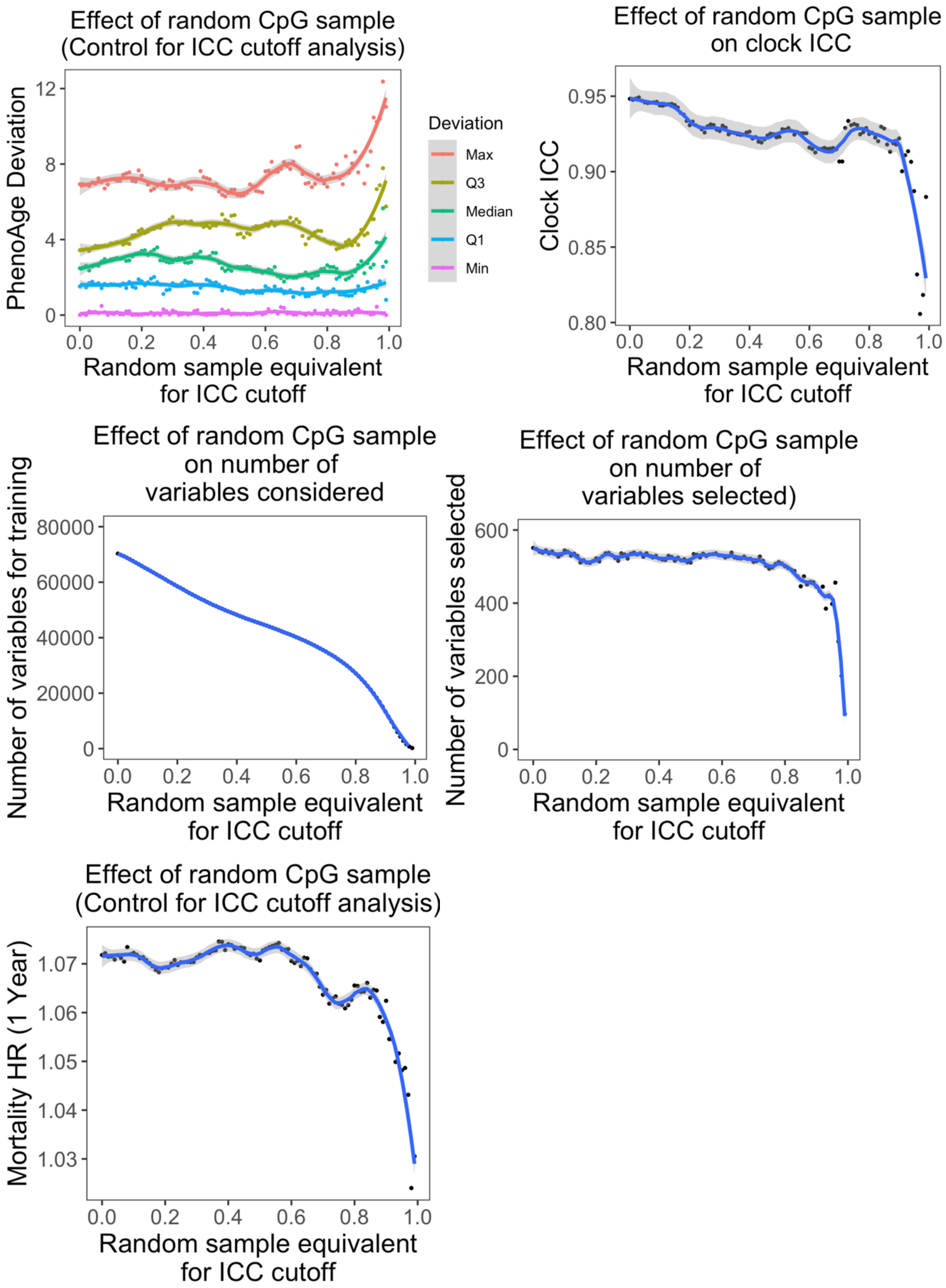

**Figure S17 CpG contributions to PC clocks.** To calculate the contribution of each CpG to the final PC clocks, we multiplied the CpG loadings for each PC by the PC weight in the clock, calculated the sum for each CpG, and divided by CpG standard deviation from the PC clock training data. CpGs present in the original clock are denoted in red.

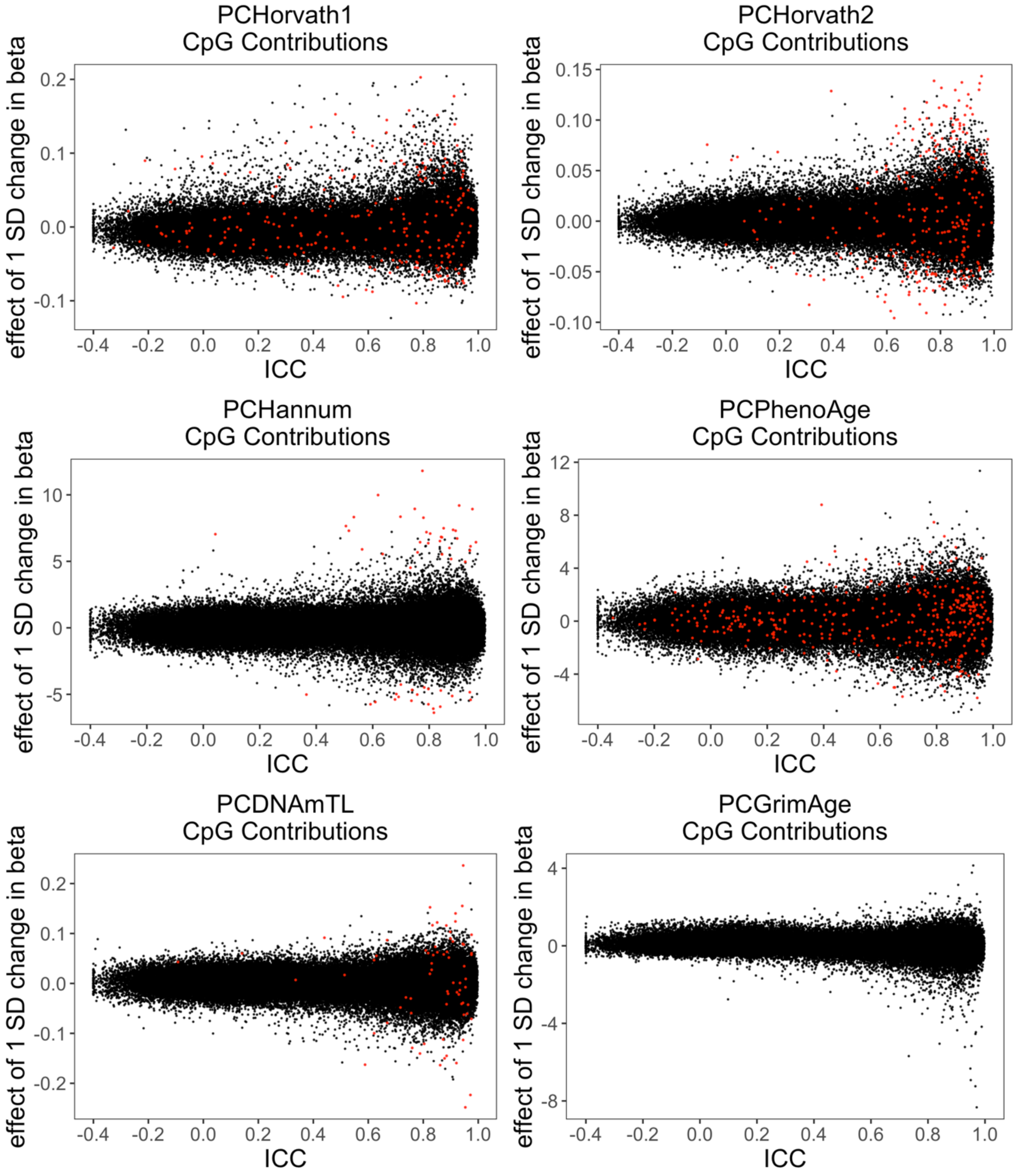

**Figure S18. Ridge plot demonstrating the distributions of clock values for cerebellum (GSE43414).**

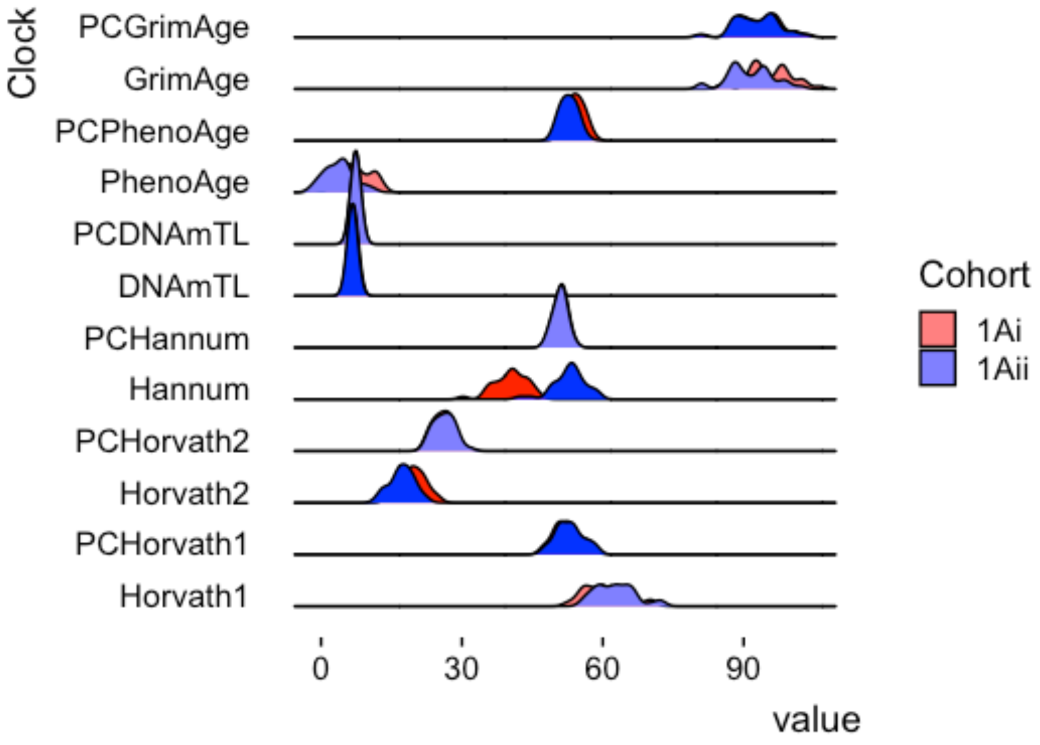

**Figure S19. Short-term longitudinal data in combat-exposed military personnel**

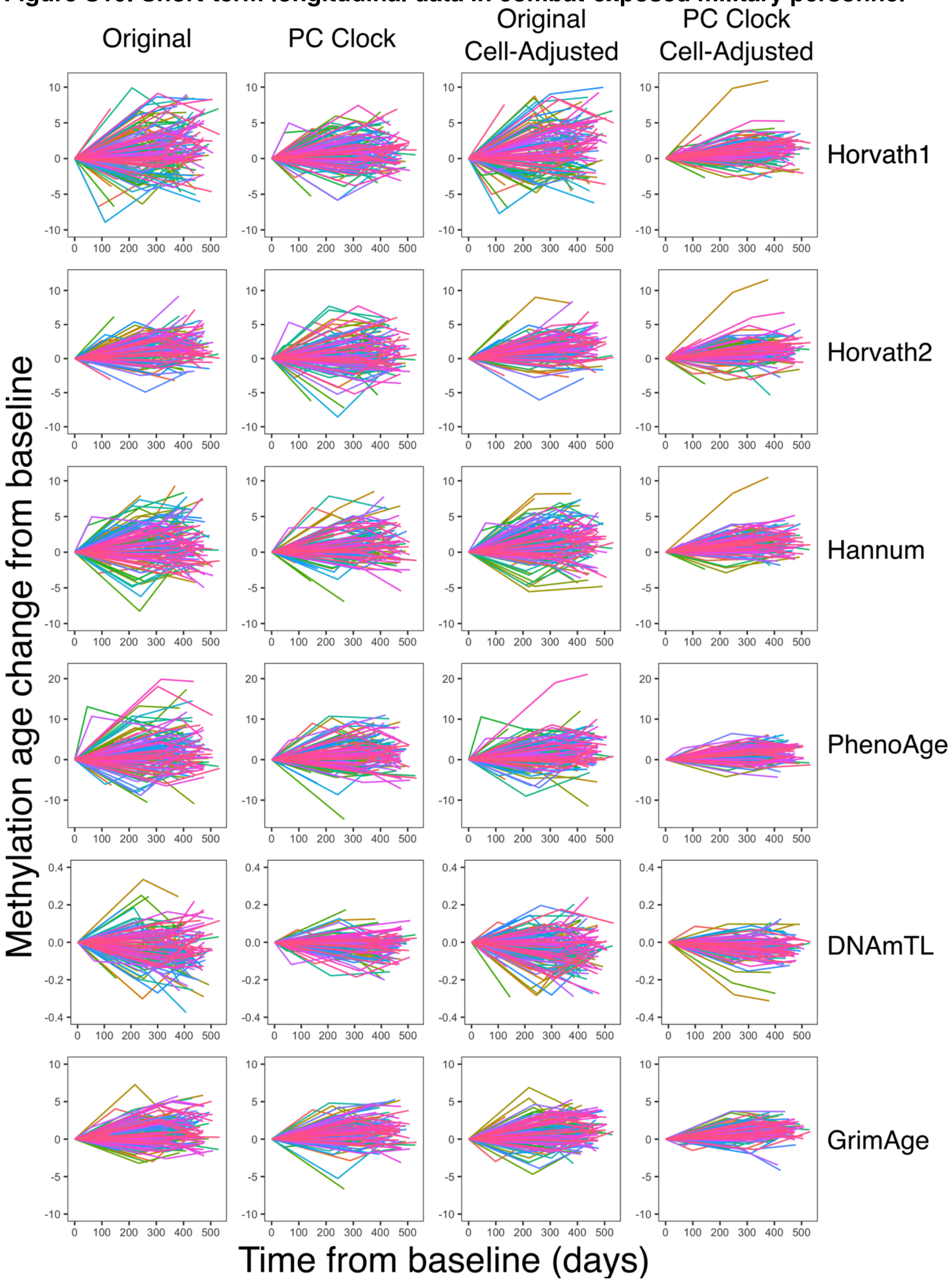

**Figure S20. Correlations between short-term longitudinal changes in clocks.** Slope was calculated for every individual for each clock, then correlated with each other.

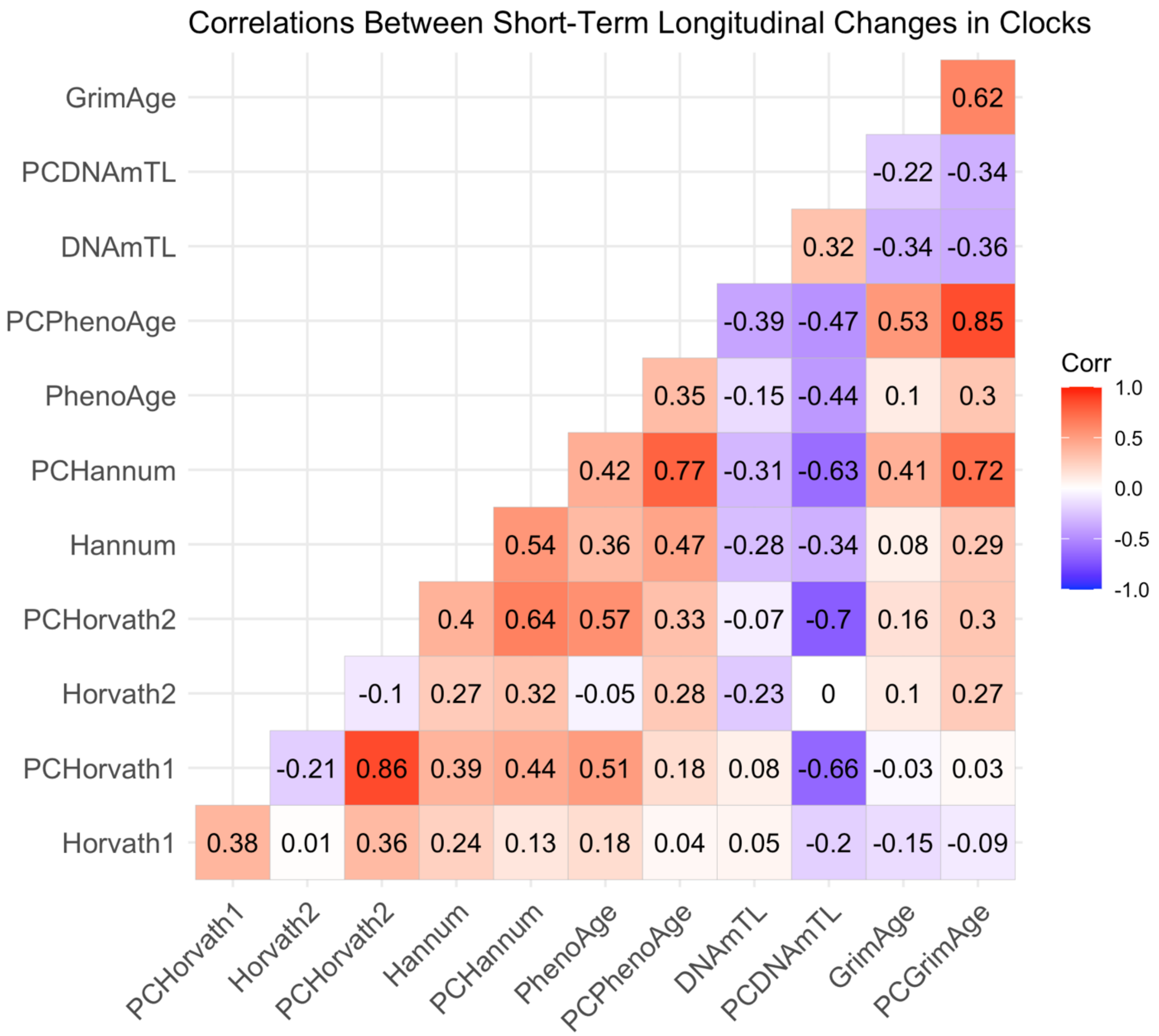

**Figure S21. Within-individual reliability of epigenetic age acceleration in short-term longitudinal data.**

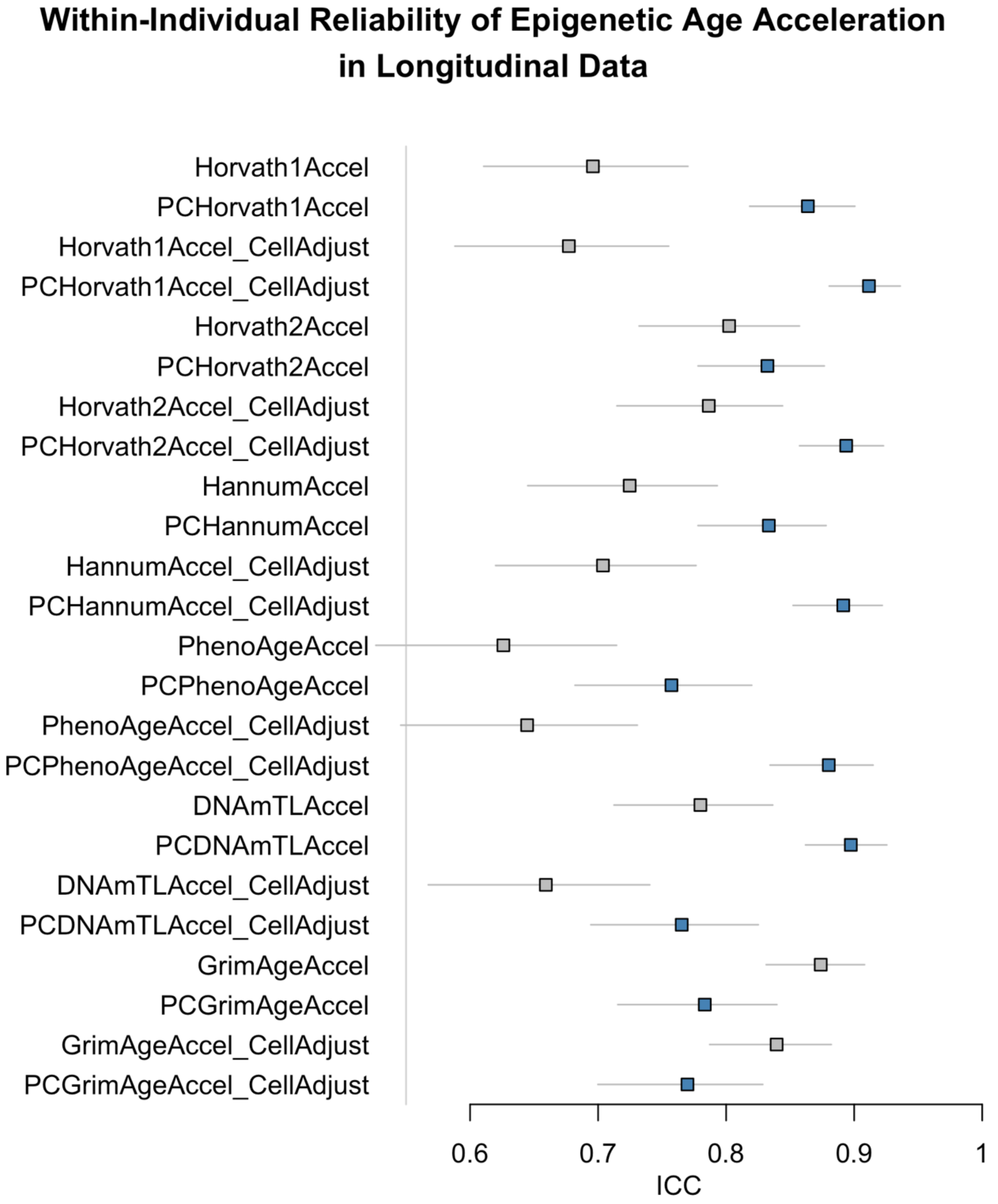

**Figure S22. Short-term longitudinal changes in epigenetic age after initiation of clozapine in schizophrenia patients.**

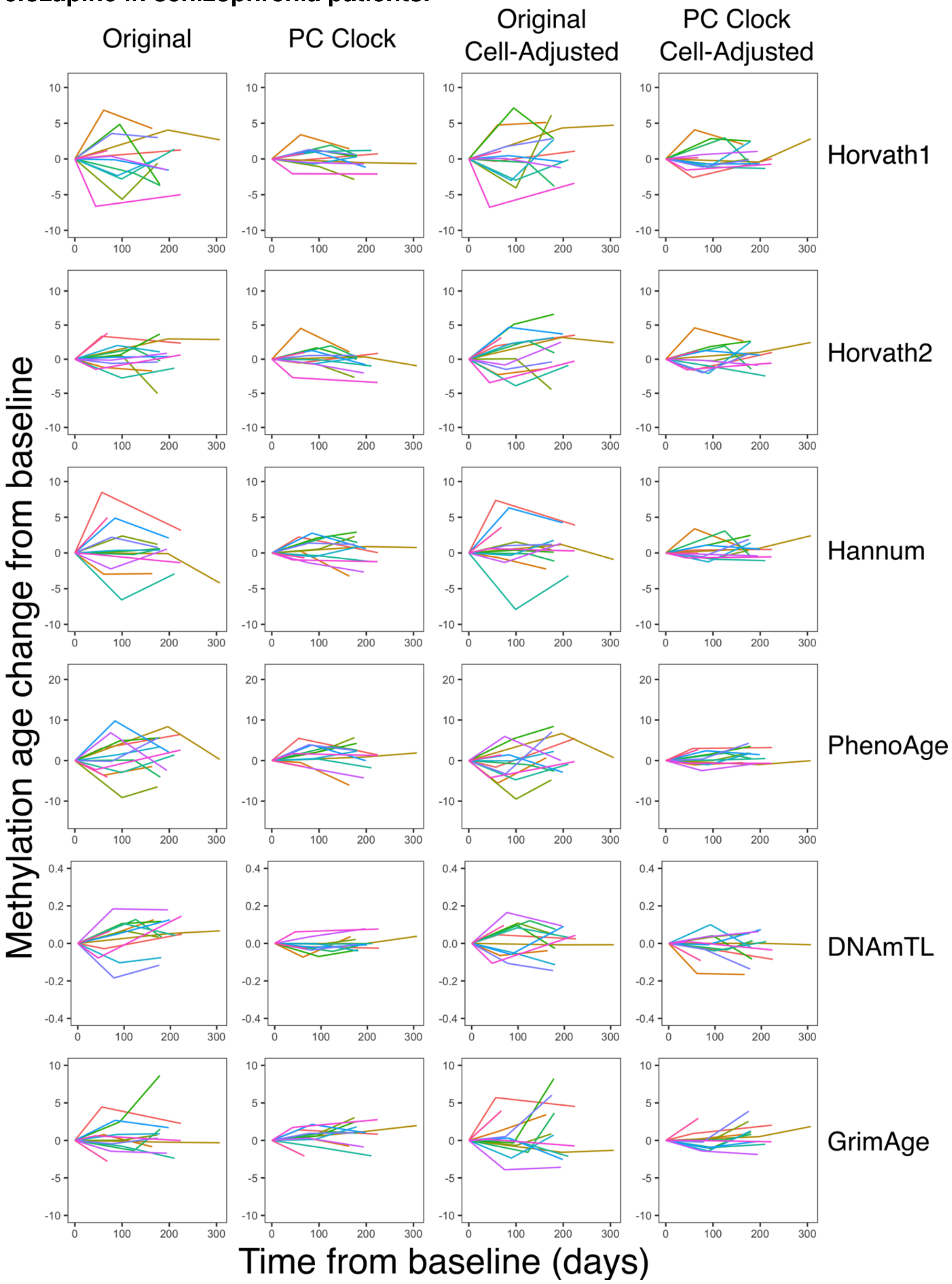

**Figure S23. Population doubling by passage in cultured primary astrocytes.**

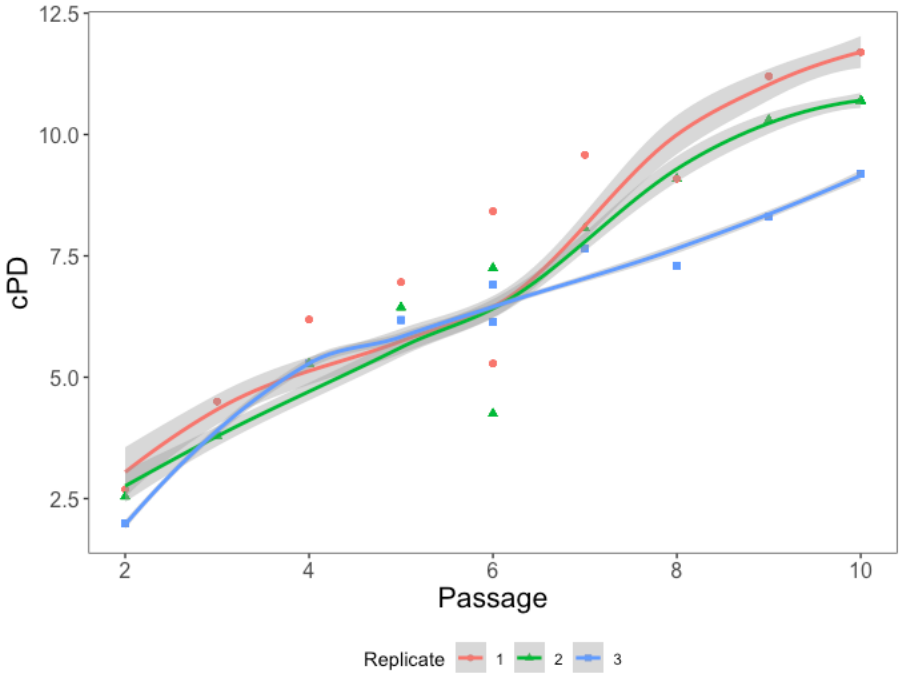
